# Metapangenomics reveals host-driven adaptations of *Methylobacterium* to the phyllosphere

**DOI:** 10.64898/2026.07.29.741493

**Authors:** Jocelyn Lauzon, Jean-Baptiste Leducq, Steven W. Kembel

**Author notes:** Corresponding author, Université du Québec à Montréal, C.P. 8888, Succ. Centre-ville, Montréal, QC H3C 3P8, Canada. **Study Funding** S.K.: Natural Sciences and Engineering Research Council of Canada (NSERC) Discovery Grant, Canada Research Chairs program; J.L.: NSERC Canada Graduate Scholarships – Master’s (CGS M), Fonds de recherche du Québec – Nature et Technologies (FRQNT) Master’s Research Scholarships; JBL: NSERC Discovery Grant.

## Abstract

Bacteria inhabiting leaf surfaces – the phyllosphere – are crucial to plant health and ecosystem functioning. *Methylobacterium* is a taxonomically diverse, growth-promoting genus, ubiquitous on leaves. Different plant species host distinct *Methylobacterium* communities, but the genomic and functional basis of *Methylobacterium* symbiotic associations with particular host species remains poorly understood. Here, we used a metapangenomic approach to quantify the influence of host species on *Methylobacterium* assemblages, to identify genes potentially involved in *Methylobacterium* adaptations to host species, and to evaluate the contribution of *Methylobacterium*’s accessory pangenome to these adaptations. We sequenced the metagenomes of 25 phyllosphere communities spanning five host species in a temperate forest in Quebec, Canada, and mapped these metagenomes onto *Methylobacterium*’s pangenome to obtain nucleotide-level coverage and composition for each population on each individual host. We revealed strong divergences in the species- and gene-level community structure of *Methylobacterium*, driven by host phylogeny and plant form. Conifer communities were notably enriched in genes involved in amino acid, lipid, and carbohydrate metabolism; broadleaves, in genes involved in cell membrane, signalling, defense, and chemotaxis; and trees, in genes related to photosynthesis, oxidative phosphorylation, and translation. The shrub *Corylus cornuta* was a reservoir of *Methylobacterium* taxonomic diversity, and harboured numerous accessory genes under positive selection. *Methylobacterium*’s accessory pangenome, evolving under weaker purifying selection, contributed importantly to gene-host associations, supporting its adaptive role. By linking genes to phyllosphere niches, our study shed light on the genetic basis of host adaptation and highlighted the crucial role of forest biodiversity in shaping microbial ecology and evolution.

## INTRODUCTION

The phyllosphere, the aerial parts of plants, harbours diverse microbial communities [1]. This habitat is characterized by nutrient scarcity, high UV exposure and rapid shifts in temperature and humidity, calling for particular adaptations in epiphytic symbionts [1–3]. The leaf bacterial microbiome supports key ecological functions related to host growth and protection and to ecosystem functioning [4,5]. Host species exert an important influence on bacterial community structure and assembly processes on tree leaves across biomes [6–9], along spatial location, season [7,10], and latitude [11,12]. The effect of hosts can partially be attributed to the unique leaf morphology, physiology and phytochemistry of each plant species – characteristics that impact microbial species assemblages [6,13,14]. There is evidence for adaptive matching between bacteria and their plant hosts [15,16], but the genomic basis of this adaptation remains poorly understood.

*Methylobacterium* (*Methylobacteriaceae*, *Hyphomicrobiales*, *Alphaproteobacteria*; [17]) is a diverse genus, estimated to include up to 104 species grouped in four clades [18]. Although *Methylobacterium* thrives in a wide range of natural and anthropogenic habitats [19], it is ubiquitous on leaf surfaces [1,3,6,10], considered as its main reservoir of diversity [18] – making this genus a model bacteria for the study of beneficial symbioses with plants. For instance, most *Methylobacterium* species are facultative methylotrophs [19,20], consuming methanol secreted by leaves [21] as a carbon and energy source, and can perform aerobic anoxygenic photosynthesis [20,22] – two traits conferring them an advantage in low chemical-energy environments [23] such as leaf surfaces. In return, *Methylobacterium* can stimulate plant growth [24,25] through the synthesis and secretion of auxins [20,26,27] and cytokinins [28,29], and can protect its host against pathogens by triggering an induced systemic resistance [30]. *Methylobacterium* community composition is impacted by host plant species [31,32], including in temperate forests [10], supporting the existence of species-specific interactions with hosts. However, no study has yet investigated which bacterial genes and their associated functions are contributing to these plant-bacteria symbiotic associations. Acquiring this knowledge on a genetically diversified, ubiquitous, and ecologically important genus such as *Methylobacterium* would strengthen our mechanistic understanding of the broad scale ecological pattern of plant host impacts on phyllosphere communities.

Pangenomes offer a valuable framework to study the ecology and evolution of highly diversified taxa [33]. The pangenome of *Methylobacterium* has been studied to link phenotypes to genes [20] and to reconstruct its phylogeny [18]. However, *Methylobacterium* core and accessory genes have not yet been functionally linked to its populations’ ecological niches. Metapangenomics, an approach that consists in mapping environmental metagenomic sequences to a taxon’s pangenome [34,35], makes it possible to uncover associations between genes of natural populations and environmental variables [34,36], such as habitat type [37], while allowing quantification of evolutionary processes such as selection [38–40]. Metapangenomics can thus reveal genes underlying niche adaptation, while avoiding biases inherent to cultivation or genome assembly from metagenomes [41,42].

This study asks how host tree species shape the ecology and evolution of *Methylobacterium* in the phyllosphere, and what are the genomic bases of bacterial adaptation to specific host species. Our first objective was to quantify the influence of host species on the diversity and composition of species and genes among *Methylobacterium* communities. Our second objective was to identify genes responsible for *Methylobacterium* adaptations to different host species, by analyzing differential gene abundances among communities, and by exploring polymorphisms in protein-coding codon sequences of *Methylobacterium* populations to detect signals of selection. Finally, since ecological niches have been hypothesized to support a local reservoir of adaptative genes that populations can acquire as accessory genes through horizontal gene transfer [43], our third objective was to evaluate the contribution of the accessory pangenome to *Methylobacterium* host species adaptations. To address these objectives, we used a metapangenomics approach: we constructed a *Methylobacterium* pangenome onto which we mapped metagenomic reads from natural phyllosphere communities sampled from five host species in a temperate forest in Quebec. We revealed strong divergences in the taxonomic and genetic community structures related to host phylogeny and plant form, and identified hundreds of potentially host-associated adaptive genes and functions belonging to both core and accessory pangenomes, linking *Methylobacterium* genes to its phyllosphere ecology.

## MATERIAL AND METHODS

### Sample collection and processing

Natural phyllosphere bacterial communities were collected at the *Station de Biologie des Laurentides* (Quebec, Canada; 45.99 N; 73.99 W) from the sub-canopy leaves of four tree species, including conifers (*Abies balsamea* and *Thuja occidentalis*) and broadleaves (*Acer saccharum* and *Fagus grandifolia*), and one broadleaf shrub species (*Corylus cornuta*). Five individuals of each species were sampled on July 26, 2023, a peak period for *Methylobacterium* diversity [10]. Sample collection, processing and DNA extraction were performed following Leducq et al. [10]. Libraries were prepared using the NEBNext Ultra II FS DNA Library Prep Kit (New England Biolabs, MA, USA), and shotgun sequencing (2x150 bp) was performed on an Illumina NovaSeq 6000 S4 flow cell using Reagent Kit v1.5. Demultiplexed paired-end reads were filtered using ‘iu-filter-quality-minoche’ function implemented in anvi’o (v8; [44,45]).

### Mapping metagenomic reads on the *Methylobacterium* pangenome

To build the *Methylobacterium* pangenome, we selected one reference genome for each of the 104 species identified in Leducq et al. [18] (Supplementary File A). For species with multiple genomes, we selected the genome with the largest N50. We performed all steps of the assembly and characterization of the *Methylobacterium* pangenome with anvi’o, with default parameters unless otherwise indicated. Within each genome, Prodigal (v2.6.3; [46]) was used (function ‘anvi-gen-contigs-database’) to predict protein-encoding genes (hereafter, simply referred to as ‘genes’) based on open reading frames (ORFs); rRNA-encoding genes were not analyzed. Amino-acid sequences of genes were functionally annotated using three databases: Clusters of Orthologous Genes (COGs; NCBI; v.20) [47], KEGG Kofam [48], and Pfam [49].

We used bowtie2 [50] to competitively map metagenomic reads from each of the 25 metagenomic samples to the 104 *Methylobacterium* genomes, then used samtools [51] to sort and index alignments. The functions ‘anvi-profile’ (--min-contig-length 1000), ‘anvi-merge’, and ‘anvi-summarize’, were used to calculate single codon variants (SCVs), horizontal coverage (i.e., coverage breath, calculated as the proportion of nucleotides having at least 1X coverage; also called ‘detection’ in anvi’o documentation) and coverage depth. For each sample, horizontal coverage and coverage depth were summarized for (i) each gene of every *Methylobacterium* species, and (ii) for each *Methylobacterium* genome.

We used the function ‘anvi-pan-genome’ (--minbit 0.5, --mcl-inflation 10), which integrates DIAMOND [52], MCL [53,54] and MUSCLE [55], to identify and align gene clusters (groups of homologous genes sharing similar amino acid sequences) and construct the pangenome. The function ‘anvi-script-compute-bayesian-pan-core’ was used to predict whether clusters corresponded to core or accessory genes [56].

### Statistical analyses

#### General approach

All statistical analyses were performed in R (v.4.4.0; [57]), using the packages tidyr [58] and dplyr [59] for data manipulation, and ggplot2 [60] and ggeffects [61] for figures. Kruskal-Wallis (KW) tests were used as a nonparametric alternative to ANOVA, with *η*^2^ as the effect size measure. All ANOVAs, KW tests and permutational multivariate analyses of variance (PERMANOVAs [62], performed with function ‘adonis2’ in vegan [63]), were followed by post-hoc tests where we compared samples 1) from *C. cornuta* to all other samples (shrub vs. trees) and 2) from *A. balsamea* and *T. occidentalis* to *A. saccharum* and *F. grandifolia* (conifer vs. broadleaf trees). Univariate post-hoc tests were made with Student’s *t*-tests, or Welch *t*-tests in the case of unequal variances. Multivariate post-hoc tests were performed by subsequent PERMANOVAs. Bonferonni correction was applied to *p*-values of all post-hoc tests. A multivariate homogeneity of variances test [64,65] (‘betadisper’, vegan) was performed after each PERMANOVA. The significance threshold was set at *α* = 0.05. Linear model assumptions were verified. Gene-level analyses were performed on gene clusters or individual genes for differential abundance or *pN*/*pS* tests, respectively; no statistical analyses were directly performed on functional annotations.

#### *Methylobacterium* within bacterial communities

To assess the relative abundance of *Methylobacterium* in whole bacterial communities, we used kraken2 (v2.1.3) [66] and bracken [67] to perform a k-mer based taxonomic annotation of all metagenomic reads and an abundance estimation at the genus level for bacteria (*Methylorubrum* reads were assigned to *Methylobacterium* [18,20]). We performed one random subsampling (‘rrarefy’, vegan) set at the lowest number of total paired-end reads of a sample (*n* = 7,597,648), then we compared *Methylobacterium* relative abundance among hosts using an ANOVA.

For beta diversity analyses, we removed any genera represented by less than 0.1% of bacterial reads in any sample, keeping 198 genera across samples (82.07% of all reads). We then performed a rarefaction (‘avgdist’, vegan; 100 iterations) set at the lowest number of total paired-end reads of a sample (*n* = 6,048,614) to obtain a rarefied Bray-Curtis distance matrix. We performed a PERMANOVA to investigate the effect of host species on the whole bacterial community composition at the genus level, and computed a Principal Coordinates Analysis (PCoA).

#### Species diversity in *Methylobacterium* communities

We assessed the detection of a species in a sample based on metagenomic read mapping onto the 104 *Methylobacterium* reference genomes, using a threshold based on outlier values of horizontal coverage. We log-transformed the coverage data for all genomes in all samples to a normal distribution (Figure S1), and calculated *z*-scores. We used a threshold of *z* ≥ 1.96 to assess outliers data points which we considered as genomes detected in a sample. The equivalent threshold in horizontal coverage was 34%.

We used the mean coverage depth (rounded to integers) of *Methylobacterium* genomes to assess species abundance in *Methylobacterium* communities. Since no correlation was found between species richness and the number of filtered reads (*p* = 0.497), we did not normalize the data prior to subsequent species-level analyses. We used ANOVAs to compare species richness and equitability (measured using the Pielou index) among hosts, and a KW test to compare Shannon index. We tested the relationship between richness and Pielou index with a linear regression. We computed a PcoA and a PERMANOVA on a Bray-Curtis distance matrix to test how the composition of *Methylobacterium* species communities varied among hosts.

To perform phylogenetic analyses on *Methylobacterium* species communities, we used a maximum-likelihood phylogenomic tree of the 104 species previously inferred [18] from concatenated alignments of core gene nucleotide sequences. Firstly, we evaluated the influence of host species on the phylogenetic relatedness of communities. We computed an interspecific phylogenetic distance matrix between *Methylobacterium* species (‘cophenetic’, stats), then calculated the standardized effect size of mean pairwise distance (SES_MPD_; ‘ses.mpd’, picante [68]) ignoring species abundance and using shuffled tip labels of a pruned tree as a null model. The obtained *z*-scores were used to compare the mean phylogenetic distance (MPD) between every pair of species within observed communities to null communities, and to compare phylogenetic clustering among host species (ANOVA). Secondly, we tested if host species harboured distinct phylogenetic groups of *Methylobacterium*. We measured the intercommunity MPD (i.e., the MPD between every pair of species from two different communities; function ‘comdist’, picante), then used this distance matrix to perform a PERMANOVA.

#### Gene clusters diversity in *Methylobacterium* communities

To assess the presence of a given individual gene (i.e., protein-encoding gene identified via the presence of an ORF) in a given sample, we used a horizontal coverage threshold of ≥ 50% and a coverage depth threshold of ≥ 0.5X. We included genes from all reference genomes, regardless of species detection, to allow the detection of genes that could have been horizontally transferred from non-detected to detected species. We summed the rounded coverage values of all individual genes belonging to the same gene cluster, resulting in a gene cluster abundances “community” table.

Prior to conducting beta diversity analyses on gene cluster composition of *Methylobacterium* communities, we performed a rarefaction on the abundances table (‘avgdist’; 44,221 counts; 100 iterations) to obtain a rarefied Bray-Curtis distance matrix (two samples were lost). We tested if cluster composition varied among host species by performing a PERMANOVA and a PCoA.

To perform alpha diversity and differential abundance analyses on gene clusters of *Methylobacterium* communities, we generated one random subsampling of the original cluster abundance table (‘rrarefy’; 44,221 counts), then filtered remaining low abundant clusters (< 0.001% of total cluster counts), resulting in 10,091 clusters left across 23 samples. We compared cluster richness and Shannon index among host species using an ANOVA and a KW test respectively. We used this rarefied and filtered table as an input to DESeq2 [69], to identify differentially abundant gene clusters (DAGCs) among host species and types. For each type of comparison, we summed the abundance of DAGCs in each COG categories and KEGG categories (manually curated based on the BRITE hierarchy, details in Table S1).

#### pN/pS ratio based on single codon variants

We analyzed codon variation in genes that mapped to genomes of the four most prevalent *Methylobacterium* species across samples – *M. sp. 018*, *M. sp. 021*, *M. sp. 022*, and *M. sp. 024 –* to detect non-synonymous and synonymous polymorphisms, as well as signals of selection. From the output of ‘anvi-profile‘, we used the functions ‘anvi-gen-variability-profile’ (--engine CDN) and ‘anvi-get-pn-ps-ratio’ (--min-departure-from-consensus 0.05 --min-coverage 10 --minimum-num-variants 10) to calculate the rate of non-synonymous polymorphism divided by the rate of synonymous polymorphism (*pN*/*pS* ratio [70]), for each gene of the four *Methylobacterium* species in each sample in which they were detected. The *pN*/*pS*^(gene)^ was calculated relatively to within sample consensus codon sequence. The *pN*/*pS* ratio indicated the direction and strength of selection; lowest ratios were interpreted as stronger negative (purifying) selection [39].

We built three linear models to identify factors influencing the *pN*/*pS* ratio per gene (*pN*/*pS*^(gene)^; response variable, log transformed). Firstly, we evaluated the effect of gene type (core or accessory), *Methylobacterium* species, host species, and all interactions, on the *pN*/*pS* ratio (Table S2a). Secondly, we evaluated the effect of genome coverage on the *pN*/*pS* ratio according to gene type and *Methylobacterium* species (Table S3a). Thirdly, we evaluated if the *pN*/*pS* ratio varied with functional categories, by adding to the first model COG category as a predictor (Table S4a). We used the package emmeans [71] to calculate estimated marginal means (EMMs) and linear trends, and to perform post-hoc tests.

To identify genes potentially evolving under positive selection in association with a particular host species, we first identified all genes that had at least one *pN*/*pS* > 1 in any sample, then we restricted analysis for genes showing codon-level variability in at least three samples from the same host. Assuming that *pN*/*pS* > 1 reflects positive selection, we calculated the mean log(*pN*/*pS*^(gene)^) across samples of the same host species, and identified all gene/host species combinations with a mean log(*pN*/*pS*^(gene)^) > 0 and a lower confidence interval > 0.

## RESULTS

### *Methylobacterium* has a large accessory pangenome

Prodigal identified 594,053 individual genes (within contigs ≥ 1000 nucleotides; excluding rRNA) in the *Methylobacterium* pangenome; COG functions were assigned to 435,527 genes, KOfam functions to 306,272 genes, and Pfam annotations to 448,027 genes. Individual genes were grouped in 70,695 gene clusters, of which 22,190 were detected in sampled communities. We identified 2214 core clusters (3.13% of the pangenome), and 68,481 accessory clusters (96.87%), of which 38,190 were singletons (Figure S2). Hypothetical proteins (i.e., gene clusters with unknown function) accounted for 108 core clusters and 37,968 accessory clusters. The pangenome showed an open pattern (methodological details in Figure S3).

### Methylobacterium was more prevalent on Corylus cornuta

A mean of 85,081,000 ± 13,449,951 (SD, standard deviation) raw paired-end reads per sample were obtained from metagenomic sequencing (Table S5). *A. saccharum* samples tended to yield fewer paired-end reads per sample (ANOVA, *p* = 0.063, adj.R^2^ = 0.217; Figure S4). The percentage of reads that mapped to the *Methylobacterium* pangenome varied among host species (KW, *p* = 0.021, *η*^2^ = 0.377) (Figure S5). Based on the kraken2/bracken taxonomic assignment, the relative abundance of *Methylobacterium* in bacterial communities also varied among host species (ANOVA, *p* < 0.001, adj.R^2^ = 0.692), with *Methylobacterium* being 2.6 times more prevalent on *C. cornuta* (mean of 13.42 ± 3.97%) than on tree species (mean of 5.11 ± 1.47%) (Figure 1; Table S6, Figure S6). Host species explained 47.0% of genera compositional variation among bacterial communities (PERMANOVA, *p* < 0.001, R^2^ = 0.471; betadisper, *p* = 0.407), and conifers harboured more homogenous communities than broadleaves (Table S7, Figure S7).

**Figure 1.**
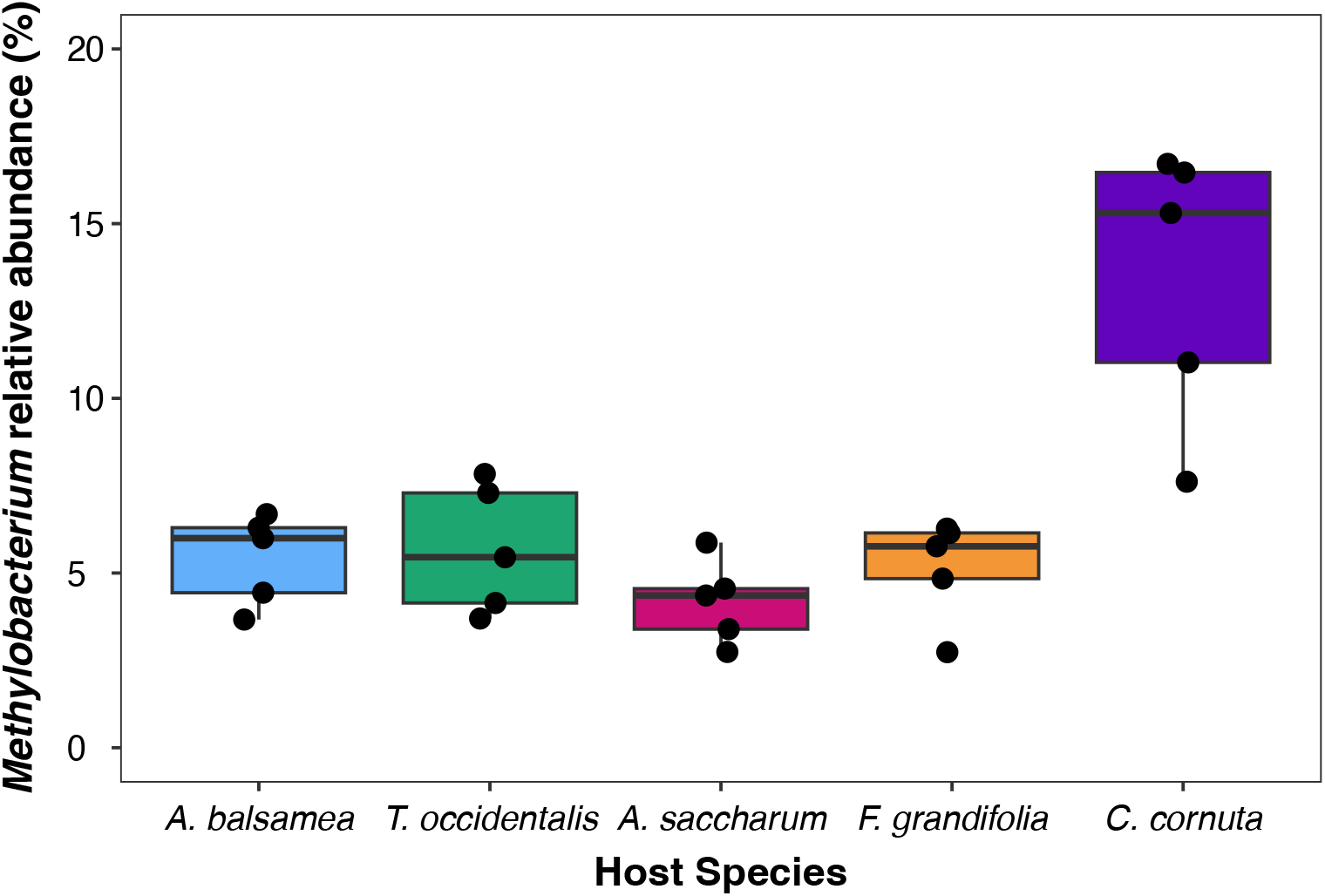
*Methylobacterium* was more prevalent on *Corylus cornuta* than on tree host species. The relative abundance of *Methylobacterium* was 2.6 times higher in *C. cornuta* phyllosphere bacterial communities (Welch *t*-test, *p* = 0.017), but did not differ between conifer and broadleaf tree species (Student *t*-test, *p* = 0.371) (Table S6). The boxplot illustrates *Methylobacterium*’s relative abundance in samples of the five host species. Across all samples, *Methylobacterium*’s relative abundance varied between 2.7 to 16.7% (mean of 6.77 ± 3.98%), and was mostly explained by host species (ANOVA, *p* < 0.001, adj.R^2^ = 0.692). Relative abundance was measured as the percentage of bacterial reads assigned to *Methylobacterium* (including *Methylorubrum*) based on a k-mer classification by kraken2/bracken. The number of reads per sample assigned to *Methylobacterium* (including *Methylorubrum*) by kraken2/bracken was correlated with the number of reads that were mapped to the *Methylobacterium* pangenome (*p* < 0.001, adj.R^2^ = 0.970).

### *Methylobacterium* taxonomic communities varied among host species

We detected a total of 23 *Methylobacterium* species in 24 samples. *Methylobacterium sp. 018*, *M. sp. 021*, *M. sp. 022*, and *M. sp. 024* were the four most abundant species in terms of total genome coverage across all hosts (Figure 2). *Methylobacterium* species richness differed among host species (ANOVA, *p* = 0.007, adj.R^2^ = 0.403) and was on average higher in *C. cornuta* leaf communities (mean of 13.6 ± 4.3 species) than on tree leaf communities (6.8 ± 3.2 species) (Table S8, Figure S8). *C. cornuta* samples also tended to harbour less even communities (Figure S9). The Pielou index decreased with species richness (linear regression, slope = –0.013, *p* = 0.004, adj.R^2^ = 0.297), indicating that rich communities were dominated by a few very abundant species.

**Figure 2.**
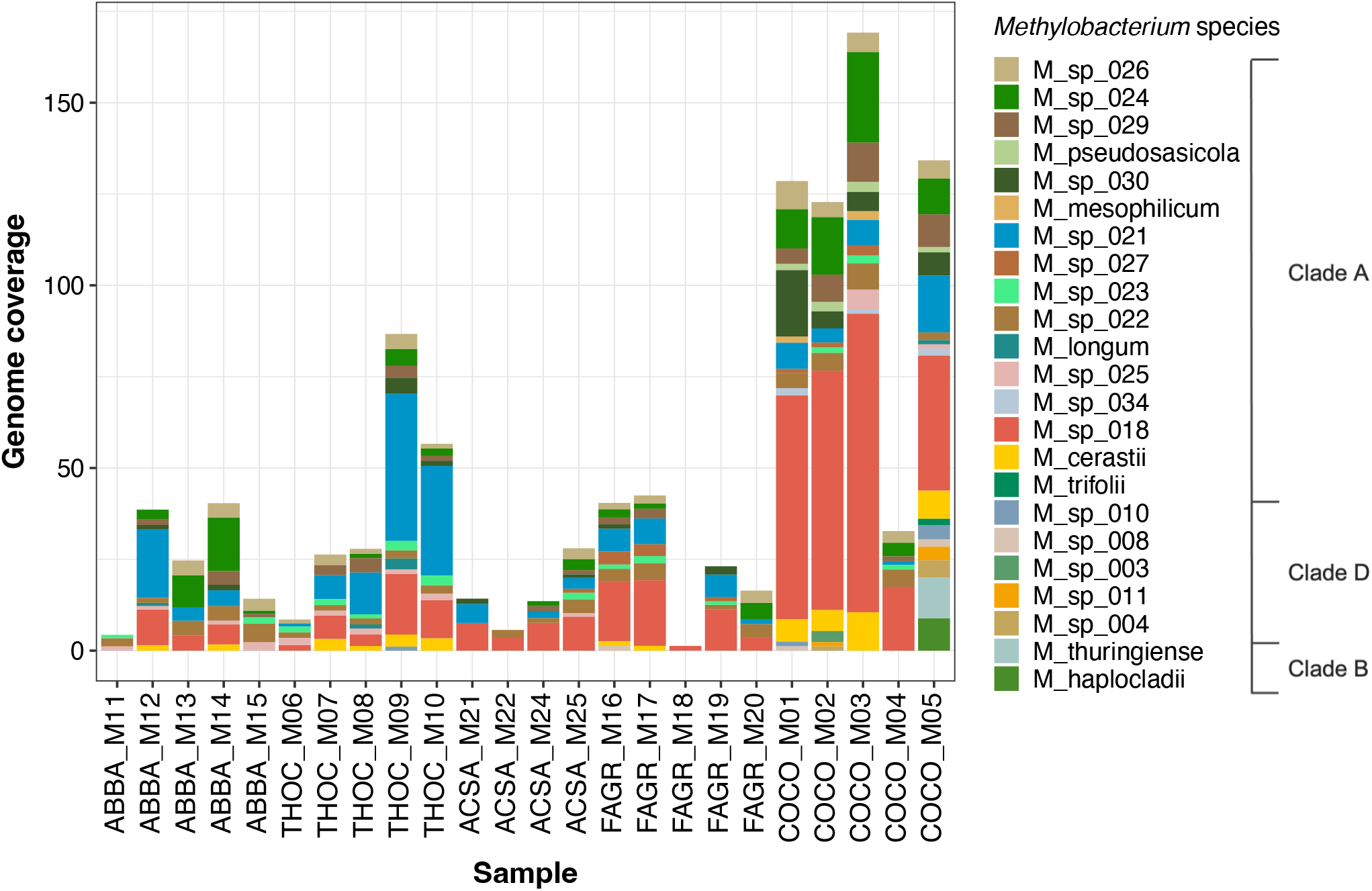
*Methylobacterium sp. 018*, *M. sp. 021*, *M. sp. 022* and *M. sp. 024* were the four most abundant species in terms of total genome coverage across all hosts. A total of 23 *Methylobacterium* species – colour-coded and listed according to their corresponding clade – were detected in 24 samples. Except for *M. cerastii*, all the Methylobacterium species that were the most prevalent in our samples are new candidate species whose strains have previously been isolated from either our study site or from another deciduous forest of southern Quebec [27]. Those strains had been isolated from either *A. saccharum* or *F. grandifolia* (see Leducq et al. [10]), but we did not observe any pattern of association between the original strain’s host species and the abundance of the corresponding Methylobacterium species on the original host. ABBA, *Abies balsamea*; ACSA, *Acer saccharum*; COCO, *Corylus cornuta*; FAGR, *Fagus grandifolia*; THOC, *Thuja occidentalis*.

Host species explained 35.5% of the compositional differences among *Methylobacterium* species communities (PERMANOVA, *p* = 0.001, R^2^ = 0.355; betadisper, *p* = 0.762). *C. cornuta* samples harboured *Methylobacterium* species communities distinct from tree species (Figure 3A). Within tree species, conifers and broadleaves differed in *Methylobacterium* species composition (Table S9).

**Figure 3.**
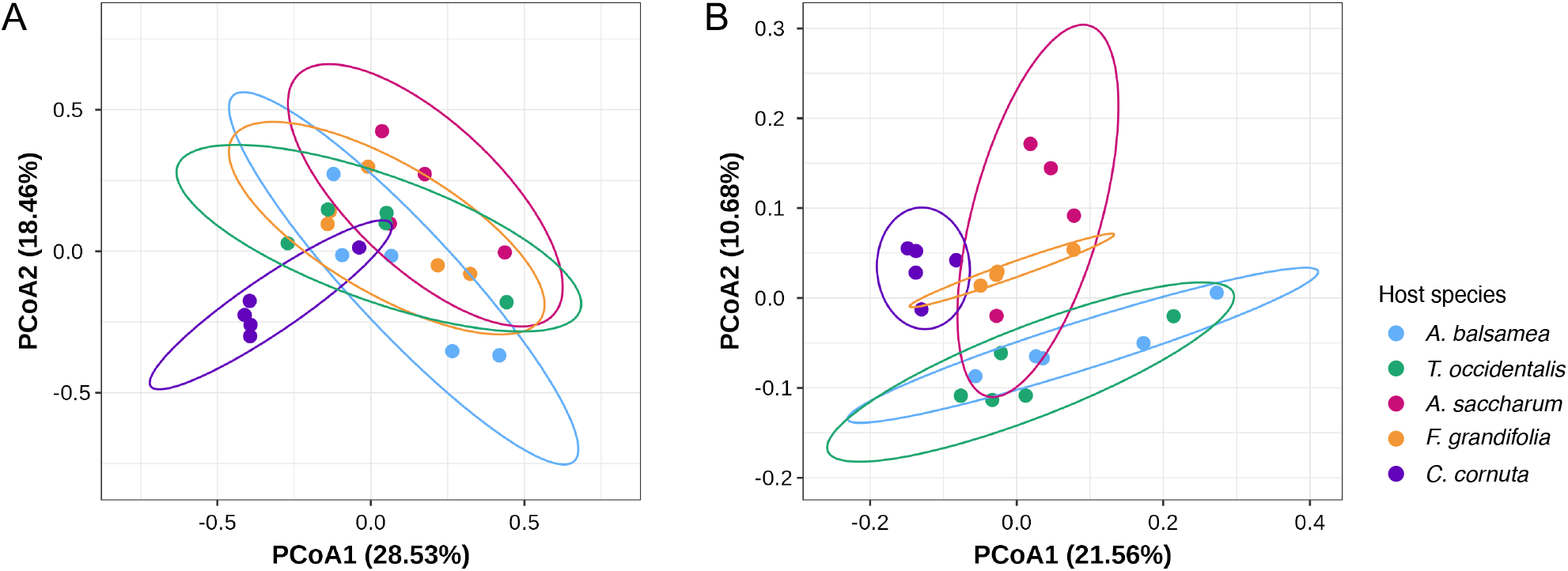
Host species impact *Methylobacterium* community composition at both species and gene levels. (A) *Methylobacterium* taxonomic community composition differed markedly between *C. cornuta* and tree species (PERMANOVA, *p* < 0.001), as well as between conifer and broadleaf trees communities (PERMANOVA, *p* = 0.048) (Table S9). (B) Gene cluster composition of *Methylobacterium* communities differed between *C. cornuta* and tree species (PERMANOVA, *p =* 0.004), as well as between conifer and broadleaf species (PERMANOVA, *p =* 0.006). Shrub-inhabiting communities had a more homogenous gene composition than tree-inhabiting communities (betadisper, *p* = 0.003) (Table S11). Bray-Curtis dissimilarity matrix was computed to calculate distances between pairs of samples. Rarefaction was performed on gene clusters counts only (*n* = 44,221 counts; 100 iterations), not on species’ genome coverage. Ellipses indicate 95% confidence intervals.

Most *Methylobacterium* communities were composed of phylogenetically closely related species (mean of MPD *z*-scores = –2.61 ± 1.41) (Table S10). Host species did not influence phylogenetic relatedness (ANOVA, *p* = 0.729, adj.R^2^ = –0.098) (Figure S10), nor did they harbour different phylogenetic groups of species (PERMANOVA, *p* = 0.215, R^2^ = 0.184) (Figure 4).

**Figure 4.**
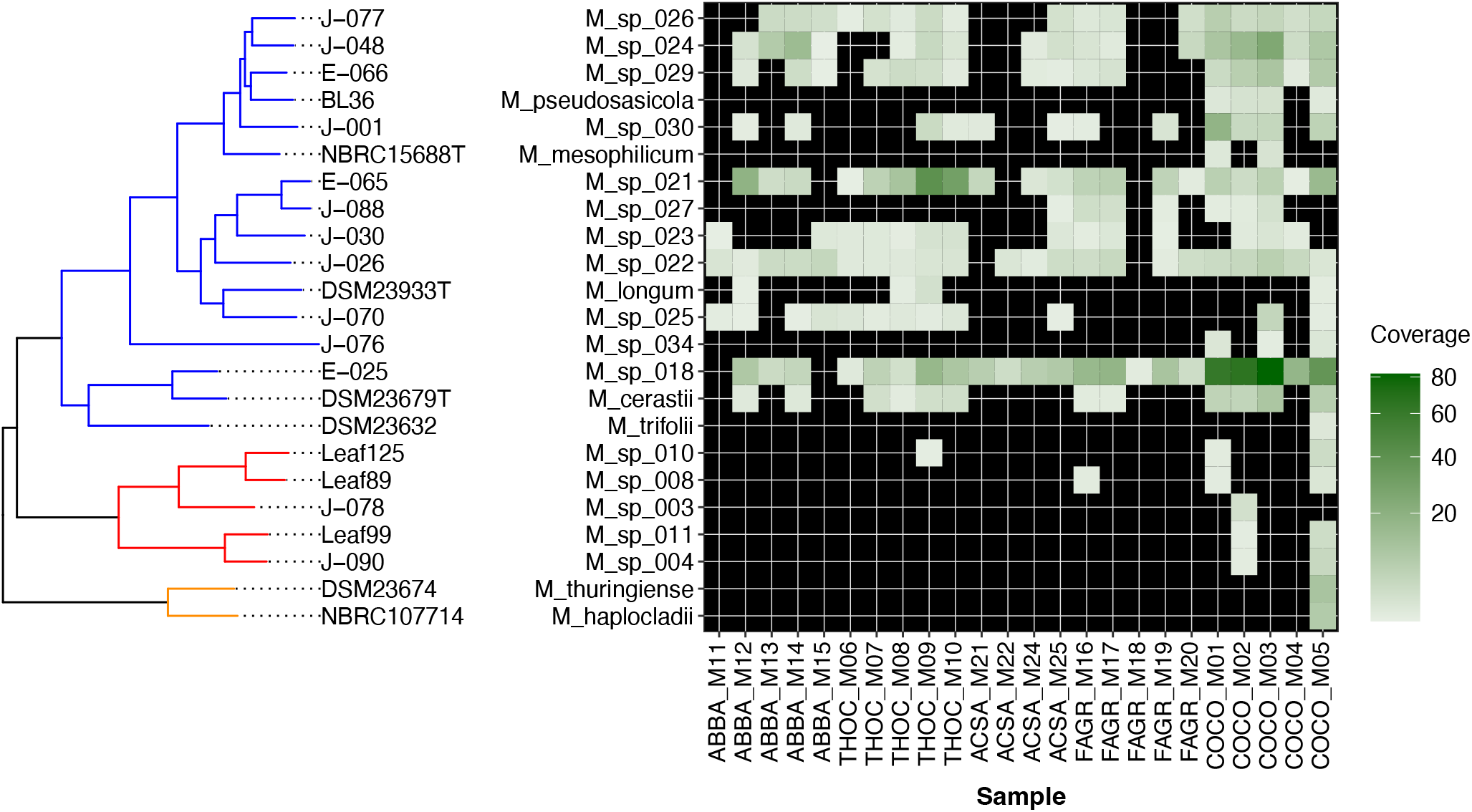
Host species did not influence the phylogenetic patterns of community assembly. Genome coverage values of the 23 detected *Methylobacterium* species, presented alongside their phylogenetic relationships, highlight the phylogenetic clustering of *Methylobacterium* communities and the dominance of clade A. Host species did not influence the degree of clustering (ANOVA, *p* = 0.729) (Figure S10) nor the phylogenetic composition of *Methylobacterium* communities (PERMANOVA, *p* = 0.215). Tree tip labels correspond to strain names whose genomes were used to represent the corresponding species. Orange-coloured tree branches represent clade B; red branches, clade D; and blue branches, clade A.

### Host species influenced the gene composition of *Methylobacterium* communities

Shifting our focus from species to genes within *Methylobacterium* communities, we found that gene cluster alpha diversity did not vary among host species (richness: ANOVA, *p* = 0.566, adj.R^2^ = – 0.046; Shannon index: KW, *p* = 0.346, *η*^2^ = 0.026). As for beta diversity, host species explained 29.6% of the variation in gene cluster composition of *Methylobacterium* communities (PERMANOVA, *p* < 0.001, R^2^ = 0.296; betadisper, *p* = 0.469). Cluster composition varied between *C. cornuta* and the tree species, and between conifers and broadleaf trees (Table S11). Cluster composition was more homogeneous among *C. cornuta* communities than among trees (Figure 3B, Table S11).

### *Methylobacterium* gene clusters were enriched according to host type

We identified 1159 gene clusters (5.22% of detected clusters) differing in abundance among host species and types. Of these differentially abundant gene clusters (DAGCs), 1050 (90.6%) were accessory clusters, and 757 (65.31%) were functionally annotated (Supplementary File B). For all comparisons between a given host species (or a group of host species) and all other species, the change in cluster abundance varied from 1.4X to 41.8X. The abundance of some clusters was driven by read recruitment from genomes of undetected species (Supplementary File C).

We identified 216 DAGCs (18.6% of all DAGCs) that differed between tree hosts and the shrub species, of which 176 were associated with trees, and 40 with *C. cornuta* (Figure 5, top panel). We found 724 DAGCs (62.5% of DAGCs) differing between broadleaf hosts (including *C. cornuta*) and conifer hosts, of which 454 were associated with broadleaves, and 270 with conifers.

**Figure 5.**
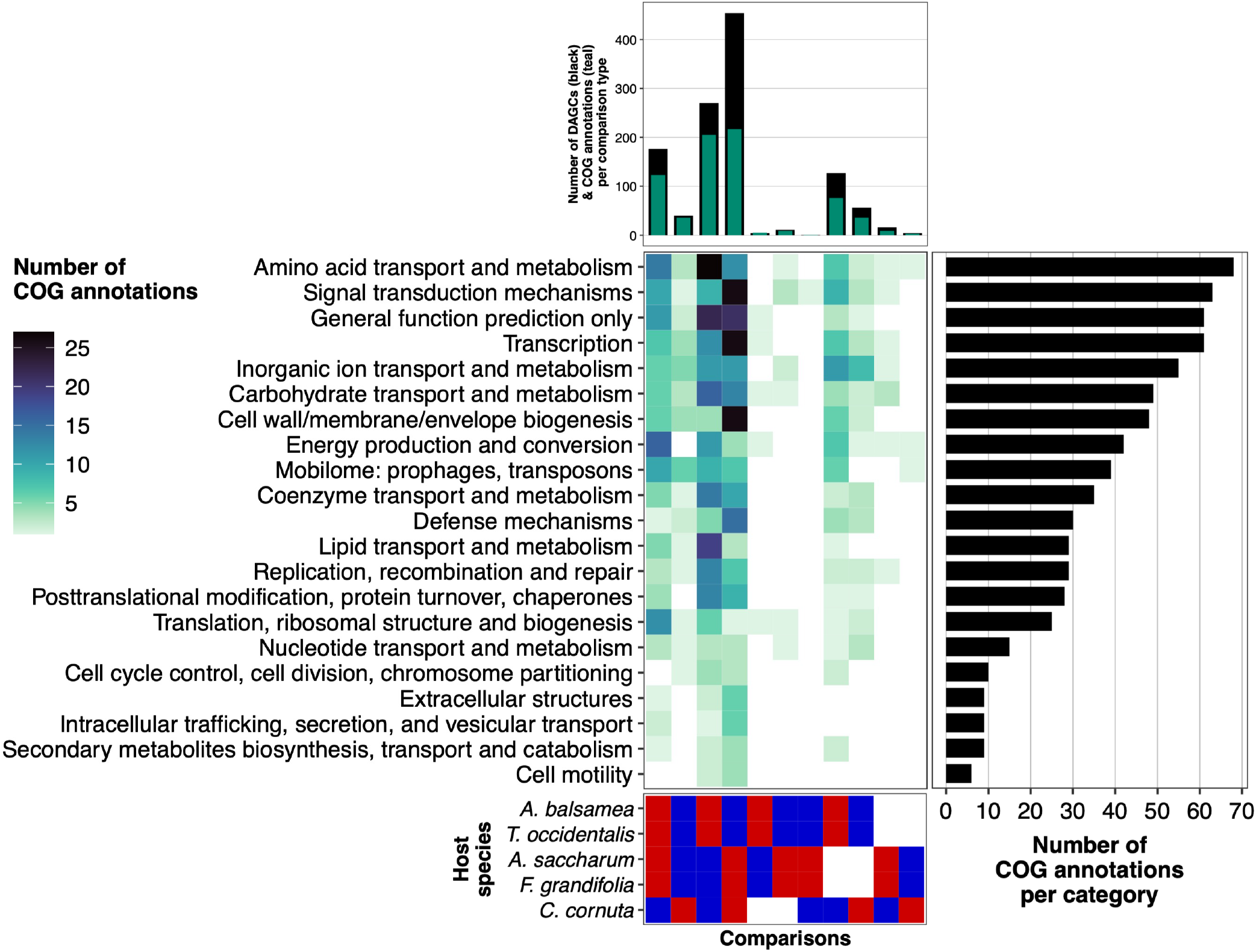
COG broad functional categories representing differentially abundant gene clusters varied greatly in proportions among hosts. The number of differentially abundant gene clusters (DAGCs) (top panel, black), as well as the abundance of COG functional categories of DAGCs (centre panel), varied among host species and host types within each comparison type. The number of COG annotations related to each category across comparison types is illustrated in the right panel: the most abundant COG categories represented across all DAGCs – accounting for more than 50% of functionally annotated DAGCs – were ‘Amino acid transport and metabolism’, ‘Signal transduction mechanisms’, ‘General function prediction only’, ‘Transcription’, ‘Inorganic ion transport and metabolism’, and ‘Carbohydrate transport and metabolism’. The number of COG annotations associated to each group of hosts across categories is illustrated in the top panel (teal). Unannotated DAGCs (*n* = 402), were only included in the calculation of the number of DAGCs per comparison (top panel, black). Only COG-annotated DAGCs (excluding the COG category ‘Function unknown’ [*n* = 47]) were used to illustrate the heatplot, and to calculate the total number of COG annotations per group of hosts (top panel, teal). DAGCs assigned to more than one COG category were counted once in each assigned category [92]. Comparison types (bottom panel) are illustrated as red and blue tiles, where red represents the focal host species or group, and blue represents the reference level against which the focal host(s) is(are) compared. No DAGC was found between the two broadleaf tree species (*A. saccharum* vs. *F. grandifolia*), nor between the two conifer species (*A. balsamea* vs. *T. occidentalis*).

We summarized functions of DAGCs using COG (Figure 5) and KEGG categories (Figure 6). Below, we report the most prevalent COG categories identified per host type, as well as distinguishing KEGG categories in parentheses. Tree-enriched DAGCs were mostly involved in energy aquisition (oxidative phosphorylation, photosynthesis, carbon fixation, nitrogen metabolism), translation (ribosome) and amino acid metabolism (alanine, aspartate, glutamate, arginine). Shrub-enriched DAGCs were related to the mobilome and inorganic ions. Conifer-enriched DAGCs were related to the transport and metabolism of amino acids (cysteine, methionine, phenylalanine, tryptophan, valine, leucine, isoleucine), of lipids (fatty acid, glycerophospholipid), and of carbohydrates (glycolysis, gluconeogenesis, propanoate), as well as in benzoate degradation (KEGG). Broadleaf-enriched DAGCs were related to cell wall and membrane (bacterial secretion system, transporters), signal transduction, transcription (transcription factors), defense (toxin-antitoxin), bacterial chemotaxis (KEGG), exopolysaccharide synthesis (KEGG) and biofilm formation (KEGG). DAGCs coding for transposases (all accessory) and ABC transporters were prevalent across all hosts.

**Figure 6.**
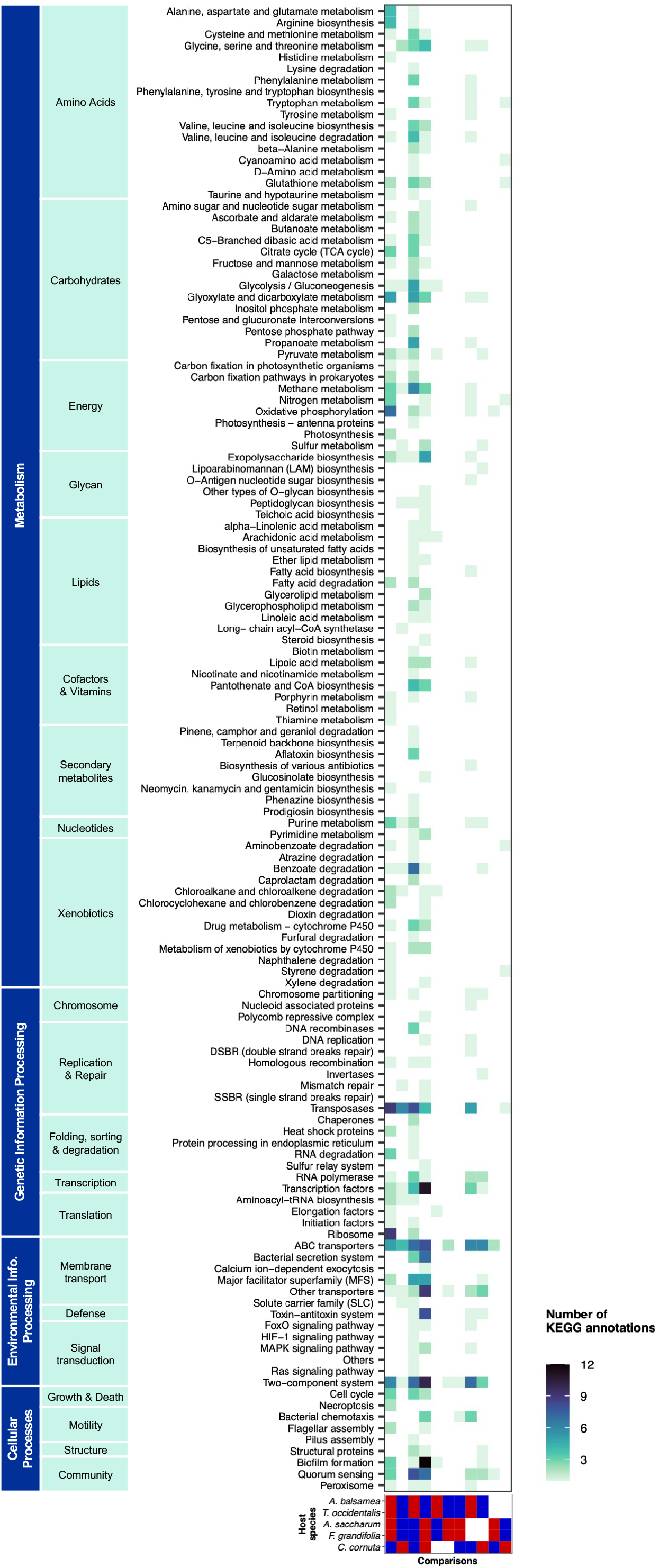
The identity and prevalence of KEGG functional categories related to differentially abundant gene clusters varied among hosts. KEGG categories were manually curated based on the BRITE hierarchy. Detailed steps performed in manual curation of KEGG categories are presented in Table S1. DAGCs assigned to more than one KEGG category were counted once in each assigned category [92]. Comparison types (bottom panel) are illustrated as red and blue tiles, where red represents the focal host species or group, and blue represents the reference level against which the focal host(s) is(are) compared. No DAGC was found between the two broadleaf tree species (*A. saccharum* vs. *F. grandifolia*), nor between the two conifer species (*A. balsamea* vs. *T. occidentalis*).

### Accessory genes showed a higher *pN*/*pS* ratio, and host-driven positive selection was more prevalent on *C. cornuta*

The *pN*/*pS*^(gene)^ ratio varied according to gene type (accessory or core), *Methylobacterium* species and host species, in interactions (linear model; *p* < 0.001; adj.R^2^ = 0.104) (Figure 7A, Table S2b). Accessory genes showed a higher *pN*/*pS* (EMM = 0.168) than core genes (0.121) across all *Methylobacterium* and host species (*p* > 0.001; Table S2c). *M. sp. 021* showed the highest *pN*/*pS*^(gene)^ across gene types and host species (EMM = 0.162), followed by *M. sp. 018*, *M. sp. 022*, and *M. sp. 024* (Table S2d). The *pN*/*pS*^(gene)^ ratio did not vary among host species when averaged across core and accessory genes and *Methylobacterium* species (Table S2e). However, regarding their accessory pangenome, conifer-inhabiting populations of *M. sp. 021* showed a higher *pN*/*pS*^(gene)^ than its broadleaf populations, while *C. cornuta* and *F. grandifolia* populations of *M. sp. 018* showed a higher *pN*/*pS*^(gene)^ than its conifer populations (Figure 7A).

**Figure 7.**
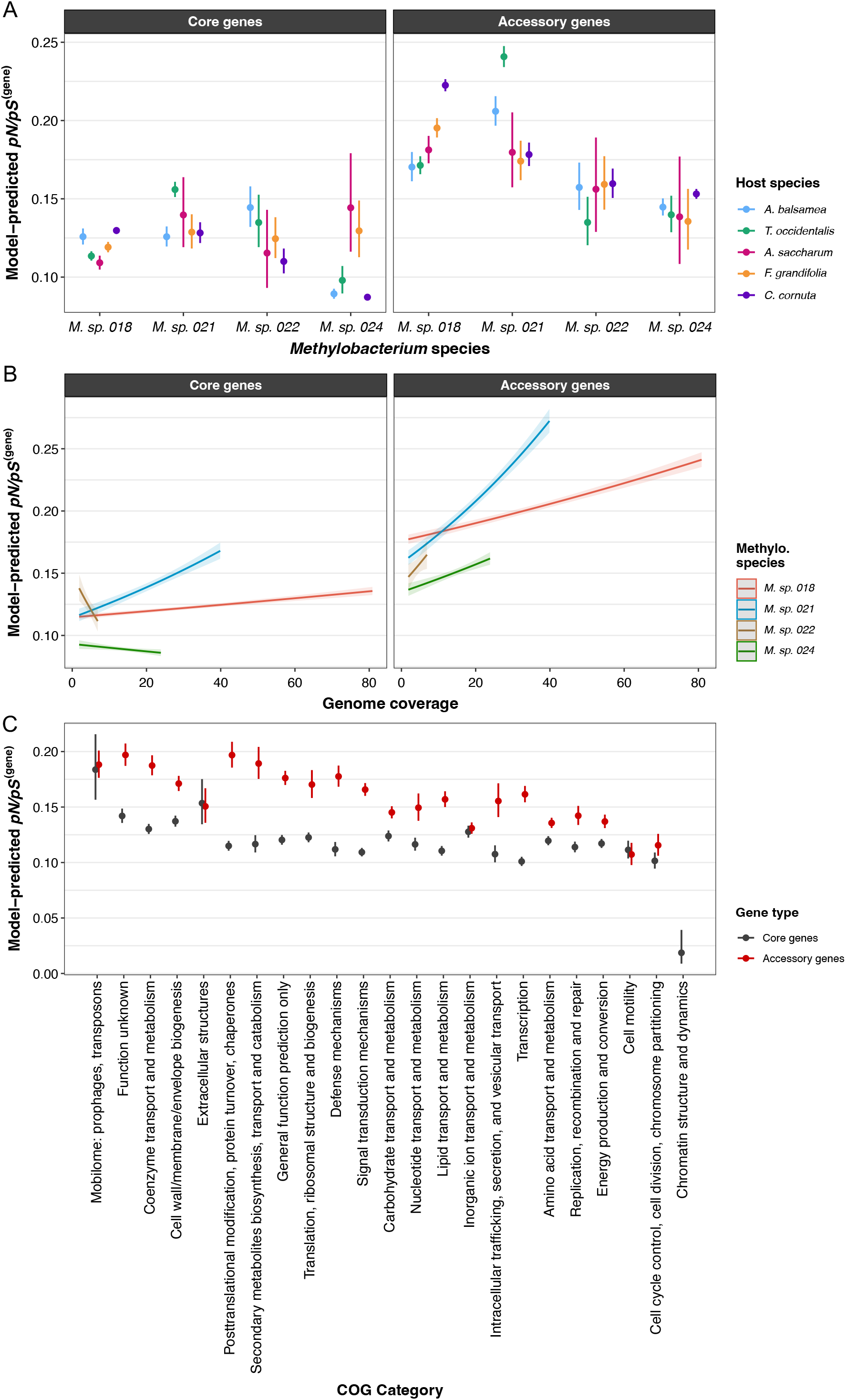
The gene-level *pN*/*pS* ratio differed according to gene type and functional category, *Methylobacterium* species and genome coverage, as well as plant host species. The *pN*/*pS* ratio represents the rate of non-synonymous polymorphism divided by the rate of synonymous polymorphism, here analyzed at the codon level of protein-encoding genes. (A) The *pN*/*pS* ratio of a gene varied depending on the interaction among its type in the pangenome (core or accessory), the *Methylobacterium* species whose genome it belongs to, and the harbouring host species. For each combination of the three factors, points represent the mean, and vertical bars, the 95% confidence interval of the mean. The linear model’s ANOVA table, estimated marginal means of each individual predictor, and pairwise comparisons among predictors’ levels are detailed in Table S2. (B) The gene-level *pN*/*pS* ratio also varied in function of the corresponding genome coverage, gene type and *Methylobacterium* species. The ranges of the illustrated predictions is limited to those of genome coverage values; lines represent the estimated marginal trends – how *pN*/*pS*^(gene)^ vary along genome coverage values –, and shaded areas, the 95% confidence interval of the mean. The linear model’s ANOVA table and the estimated marginal trends according to genome coverage, gene type (core or accessory) and *Methylobacterium* species, are presented in Table S3. (C) The gene-level *pN*/*pS* ratio was predicted for 23 COG functional categories, according to gene type (core or accessory), and averaged across all *Methylobacterium* and host species (Table S4). All alternative functional annotations of individual genes were included in the model. Categories are arranged from left to right according to decreasing estimated marginal means averaged across gene type, *Methylobacterium* species, and host species. Points represent the mean, and vertical bars, the 95% confidence interval of the mean.

The correlation between genome coverage and *pN*/*pS*^(gene)^ ratio differed according to gene type and *Methylobacterium* species (*p* < 0.001; adj.R^2^ = 0.104) (Figure 7B; Table S3b); the ratio generally increased with genome coverage, but the opposite was observed for core genes of *M. sp. 022* and *M. sp. 024* (Table S3c). No correlation was found between log(*pN*/*pS*^(gene)^) and log(gene coverage) (Pearson *r* = 0.09), excluding the possibility that gene coverage depth introduced an artifact.

The *pN*/*pS*^(gene)^ also varied by COG functional categories, in interaction with gene type (*p* < 0.001). Within the accessory pangenome, *pN*/*pS*^(gene)^ was highest for categories related to posttranslational modification, secondary metabolites, mobilome, as well as to unknown functions (Figure 7C, Table S4b). Within the core pangenome, *pN*/*pS*^(gene)^ was highest for mobilome and extracellular structures.

We found 49 unique protein-encoding genes potentially under host-associated positive selection (mean *pN*/*pS*^(gene)^ > 1; Table 1; Supplementary File D). Of those genes, 42 were identified within *M. sp. 018* population, in association with *F. grandifolia* (2 genes) or *C. cornuta* (40 genes); 6 genes within *M. sp. 021* population, in association with *T. occidentalis*; 1 gene within *M. sp. 024* population, in association with *C. cornuta*; and none in *M. sp. 022* population. All the 49 positively selected genes were part of the accessory genome (14 were functionally annotated).

**Table 1.**
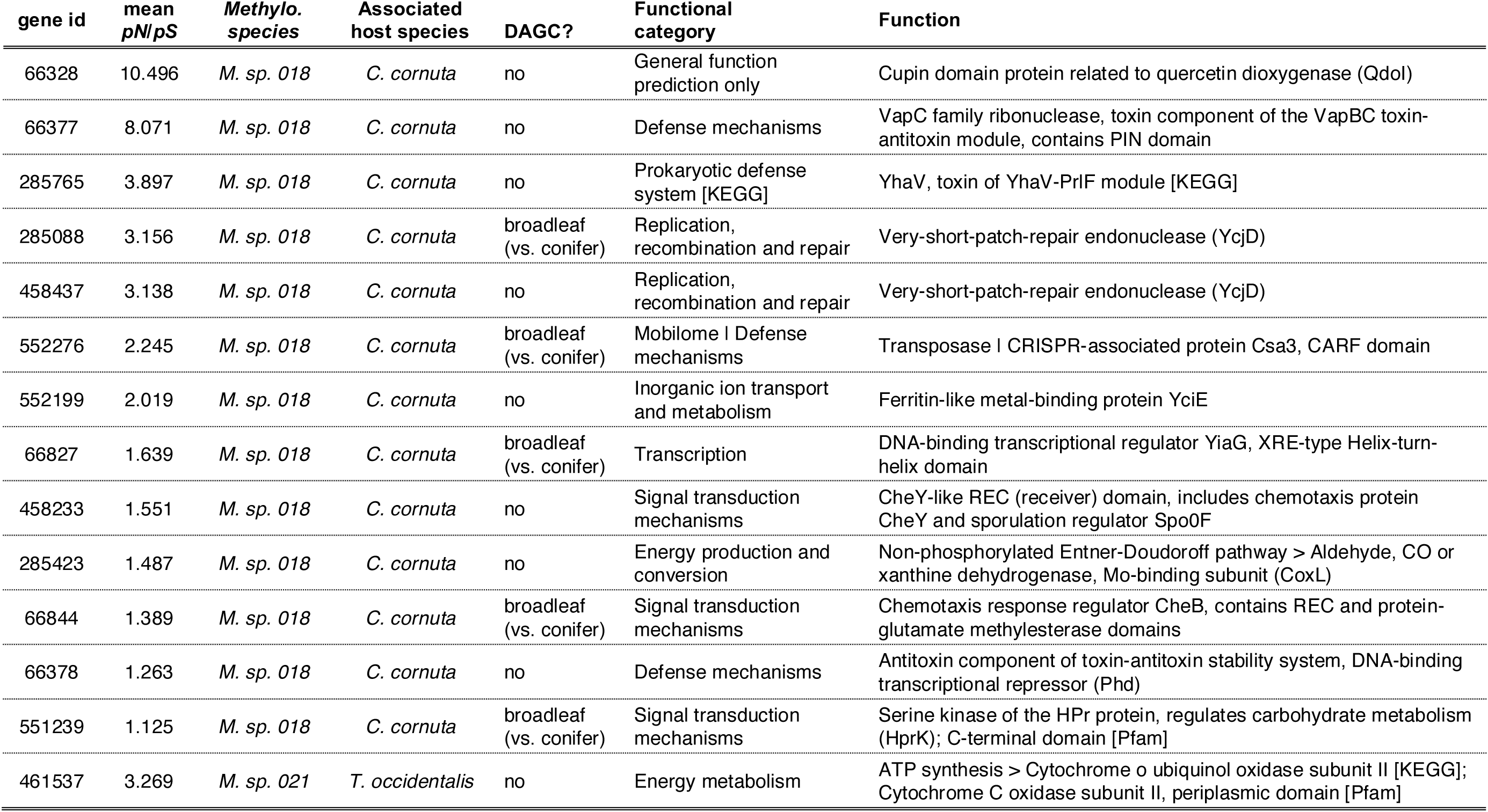
Fourteen functionally annotated accessory genes of *Methylobacterium sp. 018* were identified as being potentially under host-specific positive selection in association with *C. cornuta*, as well as one accessory gene of *M. sp. 021* in association with *T. occidentalis*. All genes presented have a mean *pN*/*pS*^(gene)^ > 1 (and a lower CI > 1) across samples of their associated host species. Functions presented in the table are inferred from COG database, unless KEGG or Pfam databases are specified in brackets. If the gene was identified as a DAGC, its comparison type is presented. The full list of potentially host-associated positively selected genes, including unannotated genes (*n* = 36) is presented in Supplementary File D.

## DISCUSSION

Our analyses demonstrated that host species is a strong driver of the taxonomic and genetic composition of *Methylobacterium* communities, in line with other studies of the tree phyllosphere [7,9,10,12,15]. With regard to our first objective, we found that host species explained around a third of the variation in *Methylobacterium* species richness (Figure S8) and in community composition at both levels of species and genes (Figure 3). Regarding our second objective, we identified hundreds of potentially host-adaptive genes and their related functions. Over 5% of gene clusters, including core and accessory clusters, were enriched in specific host types, and the proportions of functional categories representing these clusters varied greatly among hosts (Figures 5-6). Furthermore, host species influenced the strength of selection acting on some *Methylobacterium* populations, and dozens of genes were highlighted as potentially undergoing host-driven positive selection. However, hosts did not influence the phylogenetic patterns of community assembly (Figure 4), with all *Methylobacterium* communities being phylogenetically clustered, as previously observed [6,10,15].

Two recurrent patterns of divergence among host types were pervasive in all our analyses, related to host phylogeny (conifers versus broadleaves) and to plant form (trees versus shrub). Broadleaf and conifer trees harboured different *Methylobacterium* species communities (Figure 3A), as previously reported [7,15]. When considering overall bacterial genus composition, communities were more homogenous on conifers (Figure S7, Table S7), potentially reflecting leaf persistence over years. On evergreen leaves, bacterial communities could have more time to evolve into a stable state in which deterministic processes assembly processes become more important than stochastic processes [72]. In contrast, deciduous leaf communities could be more strongly shaped by massive yearly stochastic colonization events associated with early-season leaf emergence [10]. Supporting a phenology effect, conifer *Methylobacterium* communities were notably enriched in genes related to lipid metabolism (Figures 5-6), including in glycerophospholipid metabolism (e.g., phosphatidyl-N-methylethanolamine N-methyltransferase, secretory phospholipase A2), possibly favouring population survival faced with seasonal temperature variation [73,74] by modulating cell membrane properties. The fact that broadleaf communities (including *C. cornuta*) were enriched in gene clusters mostly involved in cell wall and membrane, biofilm, signalling and defense suggests adaptation to higher levels of novel biotic interactions due to more frequent colonization events. Genes involved in methyl-regulated chemotaxis (e.g., MCP, *cheR*, *cheB*, *cheY*, *cheZ*) were also enriched in broadleaves, suggesting that an enhanced motility might be more important on large leaves [15].

The shrub host *Corylus cornuta* harboured more *Methylobacterium* species than trees, and *Methylobacterium* accounted for a larger proportion of bacterial communities on *C. cornuta* compared with other hosts (Figure 1). The particularity of *C. cornuta*’s *Methylobacterium* communities could be driven by inherent leaf traits or, alternatively, by the proximity of shrub leaves to the forest soil. Soil microorganisms can colonize plant leaves [75,76], and populations can also migrate from upper strata through rainfall [77]. Vertical gradients in phyllosphere community structure, together with a higher species richness and lower evenness in the inferior strata, have previously been observed, [78,79], although the exact drivers are still not fully understood. *C. cornuta* had the highest number of host-associated positively selected genes, many of which were related to defense and signal transduction (Table 1). This could reflect a higher diversity of novel interactions due to a sustained migration, leading to an increase in positive selection [80,81] favouring genetic innovation.

Functions of enriched clusters in *Methylobacterium* communities on tree hosts (Figures 5-6) hinted at the importance of energy-related protein production and turnover. We found enriched genes involved in anoxygenic photosynthesis and oxidative phosphorylation (e.g., FoF1-type ATP synthase alpha (AtpA) and beta (AtpD) subunits, NADH-quinone oxidoreductase subunits B and I, cytochrome c oxidase subunit I, cytochrome o ubiquinol oxidase subunit I), carbon fixation or TCA cycle (e.g., malate/lactate and isocitrate dehydrogenases), and protection against heat and oxidative stresses (e.g., recombination protein RecA, chaperonins GroEL and GroES, and glucose-6-phosphate 1 involved in the pentose phosphate pathway. Notably, UV-B exposure has been found to lead to upregulation of *atpA* [82] and *recA* [83], while redirecting the carbohydrate flux from glycolysis to the pentose phosphate pathway can alleviate oxidative stress [84,85]. Tree leaves can reach higher in the canopy than shrub leaves, and thus in a closed forest receive more sunlight. Photohetereotrophic bacterial populations capable of harvesting that light more efficiently while being better able to protect themselves from heat and UV damage [86,87] would be favoured. Higher exposure to sunlight could thus be a key factor responsible for the divergence of tree and shrub-inhabiting bacterial communities in closed-canopy forest, in line with previous findings on the effect of sunlight and UV radiation [88–90]. Few gene clusters were enriched in *C. cornuta* compared to tree species indicating that adaptive genes were similar for all broadleaf host species. Although plant form (i.e., shrub or tree) exerted a strong filter on *Methylobacterium* species assemblages, host phylogeny (which is strongly correlated with leaf characteristics) might be a stronger filter than plant form in determining the phyllosphere bacterial genetic landscape.

Purifying selection was the most prevalent type of selection acting on the genes of *Methylobacterium sp. 018*, *M. sp. 021*, *M. sp. 022*, and *M. sp. 024* populations, as observed in various habitats [38–40,91]. The relatively higher *pN*/*pS*^(gene)^ ratios of *M. sp. 021* and *M. sp. 018* can be explained by a relaxed purifying selection in their large populations residing on conifers and broadleaves, respectively (Figure 7A). The general increase of the *pN*/*pS* ratio with genome coverage (Figure 7B) suggests that host species favour some specific bacterial population’s growth and diversification. COG functional categories associated with higher *pN*/*pS* ratio reflect that purifying selection is overall weaker on genes involved in environmental interaction than in basic metabolisms and cell functions (Figure 7C), corroborating other studies [70,92], and further supporting that the host habitat drive bacterial evolution.

Regarding our third objective, we found several lines of evidence that the *Methylobacterium* accessory pangenome is at least partially adaptive, in accordance with previous experimental results [93]. First, many differentially abundant accessory gene clusters recruited reads from genomes whose species were poorly covered (Supplementary File C), suggesting the possibility of adaptive horizontal gene transfer between *Methylobacterium* populations. Second, some host-enriched accessory gene clusters showed high coverage in every species that was detected in the host’s communities, suggesting that populations are maintaining these accessory genes in their genomes. Third, accessory genes had a higher *pN*/*pS* ratio than core genes, and genes potentially under positive selection were only found in the accessory pangenome. In *M. sp. 022* and *M. sp. 024*, core genes showed a decreasing *pN*/*pS* ratio with increasing population abundance, while in their accessory genes the ratio increased, suggesting different selective pressure acting on different parts of their genomes. Finally, an important fraction of host-associated positively selected accessory genes coded for proteins involved in environmental responses, thus supporting their adaptive nature. *Methylobacterium*’s pangenome was open (Figure S3), and most species were host generalists, with some populations reaching relatively large sizes. These latter observations corroborate with the hypothesis stating that large populations have large and open pangenomes because they occupy multiple niches whose selective pressures maintain accessory gene diversity [94].

In conclusion, we revealed a strong influence of host plant species on *Methylobacterium* ecology and evolution in the phyllosphere. The next step would be to broaden the range of host species, bacterial taxa, and spatial scale to look for consistent adaptive signatures to life on different hosts and to reveal generalizable eco-evolutionary patterns in the phyllosphere. Integrating plant and bacteria transcriptomics and metabolomics would also provide valuable insights into the mechanical underpinnings of host adaptations. Investigating hypothetical proteins – which accounted for a third of all DAGCs – could also reveal important niche adaptive functions [95]. Finally, processes-driven hypotheses – for example, based on leaf phenology or migration – should be considered as future experimental starting points to deepen our understanding of how forested ecosystems’ structure and biodiversity shape microbial ecology and evolution.

## Acknowledgements

We would like to thank Zihui Wang and Élanore Favron for their help in collecting and processing samples; Gabriel Lanthier for granting us access to the Station de Biologie des Laurentides (Université de Montréal); Geneviève Bourret, research professional at the Center of Excellence in Research on Orphan Diseases – Fondation Courtois (CERMO-FC) for preparing the libraries; the CHU de Québec-Université Laval Research Center for DNA sequencing; and David Ross for his support in preparing data for some analyses.

## Authors contribution

JL, SK, JBL: experimental design; JL: sample collection and processing, bioinformatic and statistical analyses, original manuscript draft; JL, SK, JBL: manuscript editing and revision.

## Data availability

Sequencing reads are available through the National Center for Biotechnology Information (NCBI) and can be found under BioSample accession numbers SAMN49978915-39. For peer-review purposes, all data and scripts have been made available through a public github repository (https://github.com/lauzonj/methylobacterium_metapangenomics).

## Supplementary Material

The ‘Supplementary Material’ document includes Supplementary Tables S1 to S11, Supplementary Figures S1 to S10, as well as textual descriptions of Supplementary Files A to D. Supplementary Files A to D can be found online at https://github.com/lauzonj/methylobacterium_metapangenomics/tree/main/supplementary_files/

