## Supplementary_Material for "Metapangenomics reveals host-driven adaptations of *Methylobacterium* to the phyllosphere"

### SUPPLEMENTARY MATERIAL (FILES, TABLES AND FIGURES)

### 1 – SUPPLEMENTARY FILES DESCRIPTIONS

**Supplementary File A** This table presents information on the 104 genomes chosen to construct *Methylobacterium's* pangenome. For the 25 genomes collected from Alessa et al. [1], the .fas files have been extracted from the .gff files contained in the article's supplementary material (as 'Data Sheet' 1 to 6), in the context of the study by Leducq et al. [2]. Clades are based on Leducq et al. [2]. genome_size_bp, number of base pairs in the genome; num_contigs, number of contigs composing the genome.

**Supplementary File B** This table presents all DAGCs functional annotations (COG, KOfam, Pfam) and differential abundance analyses' statistics (DESeq2 results). The focal host group is compared against the reference host group, the presented gene cluster is thus enriched in the focal group. type, gene type (core or accessory); genome_occurences, the number of genomes (i.e., *Methylobacterium* species) containing the gene cluster; baseMean, mean of the normalized count values; log2FoldChange, effect size; lfcSE, standard error of the log2FoldChange; padj, Benjamini–Hochberg adjusted *p*-value. '|' indicates alternative annotations originally proposed by COG or KEGG databases based on a same amino acid sequence from an individual gene; while '_or_' indicates alternative annotations when the differential abundance could arise from differentially annotated amino acid sequences from multiple individual genes.

**Supplementary File C** For every gene clusters identified as differentially abundant, this table presents: individual gene identity and its associated species genome, functional annotations (COG, KOfam, Pfam) and coverage (not normalized) for each sample. gene_callers_id, numerical identification of an individual gene (ORF) identified by Prodigal in one *Methylobcaterium* species (except for singletons, gene clusters contain multiple individual genes sharing a similar amino acid sequence); genome_name, *Methylobacterium* species; type, gene type (core or accessory); genome_occurences, the number of genomes (i.e., *Methylobacterium* species) containing the gene cluster; ABBA, *Abies balsamea*; ACSA, *Acer saccharum*; COCO, *Corylus cornuta*; FAGR, *Fagus grandifolia*; THOC, *Thuja occidentalis*.

**Supplementary File D** This table presents individual protein-coding genes potentially under positive selection in specific *Methylobacterium* species-host association, based on the $pN/pS^{(gene)}$ ratio analysis of codon polymorphism. COG, KOfam and fam functional annotations are listed. gene_callers_id, numerical identification of an individual gene (ORF) identified by Prodigal in one *Methylobacterium* species; type, gene type (core or accessory); genome_name, *Methylobacterium* species in which the gene is positively selected; host_species, host species in which the gene is positively selected (ABBA, *Abies balsamea*; ACSA, *Acer saccharum*; COCO, *Corylus cornuta*; FAGR, *Fagus grandifolia*; THOC, *Thuja occidentalis*); DAGCs, indicates whether the gene is part of a differentially abundant gene cluster and, if yes, which host group is it enriched in; genome_occurences, the number of genomes (i.e., *Methylobacterium* species) containing the gene cluster; CI_lower.log, log-scale lower bound of 95% confidence interval.

### 2 – SUPPLEMENTARY TABLES

**Table S1** Detailed steps performed in manual curation of KEGG categories, based on the BRITE hierarchy. The objective in curating the categories was to standardize category names (following ‘KEGG Orthology (KO)’ [ko00001] nomenclature), to exclude multiple annotations from similar yet different BRITE classification (duplicated information), and to exclude annotations related to general function, uncategorized or poorly characterized enzymes and proteins, as well as annotation that were uninformative in the context of our study.

| **action** | **annotations excluded (*n*)** | **DAGC removed (*n*)** |
|---|---|---|
| Excluding annotations from DAGCs that are already annotated with ‘KEGG Orthology (KO)’ (ko00001). In other words, we included annotations other than those in ‘KEGG Orthology (KO)’ only for DAGCs that were not at least once annotated as KO, to avoid duplicates). Most non-KO annotations were assigned to a relevant KO category. | 427 | 0 |
| Excluding ‘Enzymes’ (ko01000) | 19 | 0 |
| Excluding ‘Protein kinases’ (ko01001) | 4 | 0 |
| Excluding ‘Peptidases and inhibitors’ (ko01002) | 6 | 4 |
| Excluding ‘Amino acid related enzymes’ (ko01007) | 1 | 1 |
| ‘Drug metabolism – other enzymes’ (ko00001, 09100 > 09111 > 00983) | 4 | 0 |
| Excluding ‘Unclassified: metabolism > Enzymes with EC numbers’ (ko00001; 09190 > 09191 > 99980) | 27 | 27 |
| Excluding ‘Poorly characterized > General function prediction only OR Function unknown’ (ko01000; 09190 > 09194) | 24 | 24 |
| Excluding ‘Unclassified: metabolism > Others’ (ko00001; 09190 > 09191 > 99999) | 2 | 2 |
| Excluding ‘Unclassified: metabolism > Cofactor metabolism’ (ko00001; 09190 > 09191 > 99987) | 1 | 0 |
| Excluding ‘Unclassified: metabolism > Energy metabolism’ (ko00001; 09190 > 09191 > 99982) | 1 | 1 |
| Excluding ‘Unclassified: signaling and cellular processes > Others’ (ko01000; 09190 > 09193 > 99994) | 3 | 3 |
| Excluding 'Prokaryotic cytoskeleton proteins' (ko04812) | 1 | 0 |
| Excluding ‘Mitochondrial biogenesis’ (ko03029) | 4 | 1 |
| Excluding ‘Mitophagy – yeast’ (ko00001, 09140 > 09141 > 04139) | 1 | 1 |
| Excluding ‘Meiosis – yeast’ (ko00001, 09140 > 09143 > 04113) | 1 | 0 |
| Excluding ‘MAPK signaling pathway – yeast’ (ko00001, 09130 > 09132 > 04011) | 3 | 0 |
| Excluding ‘Exosome’ (ko04147) | 3 | 0 |
| **TOTAL** | **532** | **64** |

DAGC, differentially abundant gene cluster.

**Table S2** The *pN/pS*$^{(gene)}$ was predicted using a first linear model. The significance of predictors is shown in the ANOVA table, computed using sum to zero contrasts for all three factors. Estimated marginal means are presented for each predictor, averaged across the other two predictors. Pairwise comparisons were performed between gene types and *Methylobacterium* species.

a) Linear model: log(*pN/pS*$^{(gene)}$) ~ Gene type * *Methylobacterium* species * Host species

b) Type III ANOVA

| Predictor | SS | *df* | *F* | *p* |
|---|---|---|---|---|
| intercept | 44,258 | 1 | 57,753.119 | <0.001 |
| gene_type | 306 | 1 | 399.655 | <0.001 |
| meth_sp | 104 | 3 | 45.226 | <0.001 |
| host_sp | 2 | 4 | 0.603 | 0.661 |
| gene_type:meth_sp | 40 | 3 | 17.223 | <0.001 |
| gene_type:host_sp | 36 | 4 | 11.595 | <0.001 |
| meth_sp:host_sp | 264 | 12 | 39.542 | <0.001 |
| gene_type:meth_sp:host_sp | 80 | 12 | 8.697 | <0.001 |
| residuals | 57,795 | 75,418 | | |

gene_type: accessory, core; meth_sp: *M. sp. 018*, *M. sp. 021*, *M. sp. 022*, *M. sp. 024*; host_sp: *A. balsamea*, *T. occidentalis*, *A. saccharum*, *F. grandifolia*, *C. cornuta.* SS, Sum of Squares; *df*, degrees of freedom; *F,* F-statistic; *p*, *p*-value.

c) Estimated marginal means and pairwise comparisons for gene type predictor levels. Results are averaged over the levels of *Methylobacterium* species and host species.

| Gene type | EMM | SE | *df* | lower CI | upper CI |
|---|---|---|---|---|---|
| Core | 0.121 | 0.00140 | 75418 | 0.119 | 0.124 |
| Accessory | 0.168 | 0.00191 | 75418 | 0.164 | 0.172 |
| **comparison** | **ratio** | **SE** | ***df*** | ***t ratio*** | ***p*** |
| Core / Accessory | 0.723 | 0.0117 | 75418 | -19.991 | <0.0001 |

EMM, estimated marginal mean of *pN/pS*$^{(gene)}$; SE, standard error; *df*, degrees of freedom; CI, 95% confidence interval; *ratio,* proportion of EMMs difference; *p*, *p*-value.

d) Estimated marginal means and pairwise comparisons for *Methylobacterium* species predictor levels. Results are averaged over the levels of gene type and host species.

| *Methylobacterium sp.* | EMM | SE | *df* | lower CI | upper CI |
|---|---|---|---|---|---|
| *M. sp. 018* | 0.149 | 0.000864 | 75418 | 0.148 | 0.151 |
| *M. sp. 021* | 0.162 | 0.002130 | 75418 | 0.158 | 0.166 |
| *M. sp. 022* | 0.138 | 0.002820 | 75418 | 0.133 | 0.144 |
| *M. sp. 024* | 0.124 | 0.002560 | 75418 | 0.119 | 0.129 |
| **comparisons** | **ratio** | **SE** | ***df*** | ***t ratio*** | ***p*** |
| *M. sp. 018* / *M. sp. 021* | 0.922 | 0.0132 | 75418 | -5.677 | <0.0001 |
| *M. sp. 018* / *M. sp. 022* | 1.079 | 0.0228 | 75418 | 3.580 | 0.0021 |
| *M. sp. 018* / *M. sp. 024* | 1.209 | 0.0260 | 75418 | 8.824 | <0.0001 |
| *M. sp. 021* / *M. sp. 022* | 1.170 | 0.0284 | 75418 | 6.489 | <0.0001 |
| *M. sp. 021* / *M. sp. 024* | 1.312 | 0.0322 | 75418 | 11.059 | <0.0001 |
| *M. sp. 022* / *M. sp. 024* | 1.121 | 0.0326 | 75418 | 3.924 | 0.0005 |

EMM, estimated marginal mean of *pN/pS*$^{(gene)}$; SE, standard error; *df*, degrees of freedom; CI, 95% confidence interval; *ratio,* proportion of EMMs difference; *p*, *p*-value; * Bonferroni adjusted *p*-value.

e) Estimated marginal means for host species predictor levels. Results are averaged over the levels of gene type and *Methylobacterium* species.

| **Host species** | **EMM** | **SE** | ***df*** | **lower CI** | **upper CI** |
|---|---|---|---|---|---|
| *A. balsamea* | 0.142 | 0.00155 | 75418 | 0.139 | 0.145 |
| *T. occidentalis* | 0.144 | 0.00199 | 75418 | 0.140 | 0.148 |
| *A. saccharum* | 0.143 | 0.00445 | 75418 | 0.135 | 0.152 |
| *F. grandifolia* | 0.144 | 0.00253 | 75418 | 0.139 | 0.149 |
| *C. cornuta* | 0.141 | 0.00108 | 75418 | 0.139 | 0.143 |

EMM, estimated marginal mean of $pN/pS^{(gene)}$; SE, standard error; *df*, degrees of freedom; CI, 95% confidence interval.

**Table S3** The *pN/pS*$^{(gene)}$ was predicted using a second linear model. The significance of predictors is shown in the ANOVA table, computed using sum to zero contrasts for categorical factors. Estimated marginal trends are presented to show the effect of genome coverage on the *pN*/*pS* ratio, for each combination of gene type and *Methylobacterium* species.

a) Linear model: log(*pN/pS*$^{(gene)}$) ~ Gene type * *Methylobacterium* species * Genome coverage

b) Type III ANOVA

| **Predictor** | **SS** | ***df*** | ***F*** | ***p*** |
|---|---|---|---|---|
| intercept | 23,021 | 1 | 30,071.874 | <0.001 |
| gene_type | 100 | 1 | 131.156 | <0.001 |
| meth_sp | 155 | 3 | 67.6418 | <0.001 |
| genome_cov | 0 | 1 | 0.639 | 0.424 |
| gene_type:meth_sp | 32 | 3 | 14.101 | <0.001 |
| gene_type:genome_cov | 17 | 1 | 22.172 | <0.001 |
| meth_sp:genome_cov | 147 | 3 | 63.839 | <0.001 |
| gene_type:meth_sp:genome_cov | 29 | 3 | 12.461 | <0.001 |
| residuals | 57,752 | 75,442 | | |

gene_type: accessory, core; meth_sp: *M. sp. 018*, *M. sp. 021*, *M. sp. 022*, *M. sp. 024.* SS, Sum of Squares; *df*, degrees of freedom; *F,* F-statistic; *p*, *p*-value.

c) Estimated marginal trends representing the relation between *pN/pS*$^{(gene)}$ and genome coverage, according to gene type and *Methylobacterium* species. Linear trends represent how the predicted value of *pN*/*pS*$^{(gene)}$ varies with a 1X increase in genome coverage.

| **Gene type** | ***Methylobacterium* species** | **linear trend** | **SE** | ***df*** | **lower CI** | **upper CI** |
|---|---|---|---|---|---|---|
| Core | *M. sp. 018* | 0.00209 | 0.00022 | 75,442 | 0.00165 | 0.00253 |
| Core | *M. sp. 021* | 0.00966 | 0.00092 | 75,442 | 0.00785 | 0.01146 |
| Core | *M. sp. 022* | -0.04247 | 0.01270 | 75,442 | -0.0674 | -0.01757 |
| Core | *M. sp. 024* | -0.00332 | 0.00137 | 75,442 | -0.00601 | -0.00063 |
| Accessory | *M. sp. 018* | 0.00390 | 0.00024 | 75,442 | 0.00342 | 0.00438 |
| Accessory | *M. sp. 021* | 0.01360 | 0.00080 | 75,442 | 0.0120 | 0.01516 |
| Accessory | *M. sp. 022* | 0.02284 | 0.01170 | 75,442 | -0.00006 | 0.04574 |
| Accessory | *M. sp. 024* | 0.00763 | 0.00135 | 75,442 | 0.00497 | 0.01028 |

SE, standard error; *df*, degrees of freedom; CI, 95% confidence interval.

**Table S4** Comparison of model-predicted *pN*/*pS*$^{(gene)}$ among COG functional categories for core and accessory genes. All alternative functional annotations of individual genes were included to compute the model.

a) Linear model: log(*pN/pS*$^{(gene)}$) ~ Gene type * *Methylobacterium* species * Host species + COG category + Gene type:COG category

b) Estimated marginal means of *pN*/*pS*$^{(gene)}$ and 95% confidence intervals for core and accessory genes of 23 COG functional categories. Category order is based on decreasing EMM averaged across gene type, *Methylobacterium* species, and host species. Results presented by gene type are averaged over the levels of *Methylobacterium* species and host species.

| | **Core genes** | | | **Accessory genes** | | |
|---|---|---|---|---|---|---|
| **COG functional category** | **EMM** | **lower CI** | **upper CI** | **EMM** | **lower CI** | **upper CI** |
| Mobilome: prophages, transposons | 0.184 | 0.157 | 0.216 | 0.188 | 0.176 | 0.201 |
| Function unknown | 0.142 | 0.136 | 0.149 | 0.197 | 0.187 | 0.207 |
| Coenzyme transport and metabolism | 0.130 | 0.126 | 0.135 | 0.187 | 0.179 | 0.197 |
| Cell wall/membrane/envelope biogenesis | 0.137 | 0.132 | 0.142 | 0.171 | 0.164 | 0.178 |
| Extracellular structures | 0.153 | 0.134 | 0.175 | 0.151 | 0.136 | 0.167 |
| Posttranslational modification, protein turnover, chaperones | 0.115 | 0.111 | 0.120 | 0.197 | 0.186 | 0.209 |
| Secondary metabolites biosynthesis, transport and catabolism | 0.117 | 0.109 | 0.125 | 0.189 | 0.175 | 0.204 |
| General function prediction only | 0.120 | 0.116 | 0.125 | 0.176 | 0.170 | 0.183 |
| Translation, ribosomal structure and biogenesis | 0.123 | 0.118 | 0.127 | 0.170 | 0.158 | 0.183 |
| Defense mechanisms | 0.112 | 0.106 | 0.119 | 0.178 | 0.168 | 0.187 |
| Signal transduction mechanisms | 0.109 | 0.106 | 0.113 | 0.166 | 0.160 | 0.172 |
| Carbohydrate transport and metabolism | 0.124 | 0.119 | 0.129 | 0.145 | 0.140 | 0.151 |
| Nucleotide transport and metabolism | 0.116 | 0.111 | 0.122 | 0.149 | 0.138 | 0.162 |
| Lipid transport and metabolism | 0.110 | 0.106 | 0.115 | 0.157 | 0.150 | 0.164 |
| Inorganic ion transport and metabolism | 0.128 | 0.122 | 0.133 | 0.131 | 0.126 | 0.136 |
| Intracellular trafficking, secretion, and vesicular transport | 0.108 | 0.100 | 0.115 | 0.155 | 0.141 | 0.171 |
| Transcription | 0.101 | 0.097 | 0.105 | 0.161 | 0.154 | 0.169 |
| Amino acid transport and metabolism | 0.120 | 0.116 | 0.124 | 0.136 | 0.131 | 0.140 |
| Replication, recombination and repair | 0.114 | 0.109 | 0.119 | 0.142 | 0.134 | 0.151 |
| Energy production and conversion | 0.117 | 0.113 | 0.121 | 0.137 | 0.131 | 0.143 |
| Cell motility | 0.111 | 0.104 | 0.120 | 0.107 | 0.098 | 0.118 |
| Cell cycle control, cell division, chromosome partitioning | 0.102 | 0.094 | 0.109 | 0.116 | 0.106 | 0.126 |
| Chromatin structure and dynamics | 0.019 | 0.009 | 0.039 | - | - | - |

EMM, estimated marginal mean of *pN*/*pS*$^{(gene)}$; CI, 95% confidence interval.

**Table S5** Number of metagenomic paired-end reads (raw and quality filtered) obtained from sequencing, and number and percentage of reads that mapped to *the Methylobacterium* pangenome. The amount of filtered read pairs obtained showed a tendency to vary with host species (ANOVA, $p = 0.063$, adj.$R^2 = 0.217$), due to *A. saccharum* samples generally yielding fewer pairs (Figure S4). The amount of filtered single reads that were mapped to the Methylobacterium pangenome varied with host species (KW, $p = 0.015$, $\eta^2 = 0.420$), with C. cornuta samples mapping a larger proportion of Methylobacterium reads than the other four host species (Figure S5). Overall, there was no significant correlation between the number of mapped reads and the amount of filtered read pairs (regression, $p = 0.112$, Peason $r = 0.326$), and Methylobacterium species richness (Figure S8) was not correlated with the amount of filtered read pairs per sample (regression, $p = 0.497$, adj.$R^2 = -0.023$), thus indicating that our sequencing depth was large enough to capture most of Methylobacterium genomic DNA and diversity in our samples.

| sample | raw pairs | filtered pairs | filtered pairs (%) | mapped reads | mapped reads (%) |
|---|---|---|---|---|---|
| ABBA_M11 | 77366671 | 75813593 | 97,99 | 649356 | 0,42825829 |
| ABBA_M12 | 87141269 | 85384508 | 97,98 | 2278952 | 1,33452312 |
| ABBA_M13 | 88780323 | 86880987 | 97,86 | 1806157 | 1,03944319 |
| ABBA_M14 | 123574867 | 121167854 | 98,05 | 2718638 | 1,12184788 |
| ABBA_M15 | 81172304 | 79497890 | 97,94 | 1169192 | 0,7353604 |
| ACSA_M21 | 67857059 | 66571885 | 98,11 | 1248284 | 0,93754593 |
| ACSA_M22 | 82728655 | 81098799 | 98,03 | 598867 | 0,36922063 |
| ACSA_M23 | 54301257 | 53372573 | 98,29 | 275189 | 0,25780001 |
| ACSA_M24 | 64832058 | 63664639 | 98,2 | 989884 | 0,77742057 |
| ACSA_M25 | 83924385 | 82203757 | 97,95 | 1621315 | 0,98615627 |
| COCO_M01 | 75942855 | 74412420 | 97,98 | 6193356 | 4,16150691 |
| COCO_M02 | 76928927 | 75377338 | 97,98 | 5915567 | 3,92396916 |
| COCO_M03 | 87544660 | 85697936 | 97,89 | 7938674 | 4,63177666 |
| COCO_M04 | 84214348 | 82424196 | 97,87 | 1813444 | 1,10006775 |
| COCO_M05 | 92352501 | 90309976 | 97,79 | 6246367 | 3,45829291 |
| FAGR_M16 | 93003886 | 91137163 | 97,99 | 2251260 | 1,2350944 |
| FAGR_M17 | 95570792 | 93755464 | 98,1 | 2420157 | 1,29067518 |
| FAGR_M18 | 105350083 | 103332477 | 98,08 | 489432 | 0,2368239 |
| FAGR_M19 | 90247775 | 88512584 | 98,08 | 1540790 | 0,87037906 |
| FAGR_M20 | 72907395 | 71672261 | 98,31 | 1213583 | 0,84661973 |
| THOC_M06 | 85461053 | 83664129 | 97,9 | 863490 | 0,51604553 |
| THOC_M07 | 88637669 | 86818511 | 97,95 | 1669054 | 0,96123164 |
| THOC_M08 | 84190622 | 82383296 | 97,85 | 1809544 | 1,09824691 |
| THOC_M09 | 96083679 | 94188691 | 98,03 | 4635647 | 2,46082993 |
| THOC_M10 | 86909907 | 85020898 | 97,83 | 3352641 | 1,97165702 |

ABBA, *Abies balsamea*; ACSA, *Acer saccharum*; COCO, *Corylus cornuta*; FAGR, *Fagus grandifolia*; THOC, *Thuja occidentalis*.

**Table S6** Comparison of *Methylobacterium* relative abundance in bacterial communities between *C. cornuta* and all four tree host species (Welch *t*-test), and between conifer and broadleaf trees (Student *t*-test).

| comparison | *df* | *t* | *p* |
|---|---|---|---|
| shrub – trees | 4.2772 | 4.6028 | 0.01706 |
| conifer trees – broadleaf trees | 18 | 1.376 | 0.37140 |

*df*, degrees of freedom; *t*, t-statistic; *p*, *p*–value (Bonferroni adjusted).

**Table S7** Comparison of bacterial genera community composition between *C. cornuta* and all four tree host species, and between conifer and broadleaf trees, using PERMANOVAs and multivariate homogeneity of variances tests (betadisper).

| | PERMANOVA | | | | betadisper | | |
|---|---|---|---|---|---|---|---|
| ***comparison*** | ***df*** | ***F*** | ***p**** | **$R^2$** | ***df*** | ***F*** | ***p*** |
| shrub – trees | 1 | 7.4635 | 0.001 | 0.245 | 1 | 0.2004 | 0.6586 |
| conifer trees – broadleaf trees | 1 | 5.4734 | 0.001 | 0.2332 | 1 | 8.0953 | 0.01074 |

*df*, degrees of freedom; *F,* F-statistic; *p*, *p*-value; * Bonferroni adjusted *p*-value.

**Table S8** Comparison of *Methylobacterium* species richness between *C. cornuta* and all four tree host species, and between conifer and broadleaf trees, using Welch t-tests.

| comparison | *df* | *t* | *p* |
|---|---|---|---|
| shrub – trees | 5.2691 | 3.2931 | 0.0401 |
| conifer trees – broadleaf trees | 15.006 | 1.3754 | 0.3784 |

*df*, degrees of freedom; *t*, t-statistic; *p*, p–value (Bonferroni adjusted).

**Table S9** Comparison of *Methylobacterium* species community composition between *C. cornuta* and all four tree host species, and between conifer and broadleaf trees, using PERMANOVAs and multivariate homogeneity of variances tests (betadisper).

| | PERMANOVA | | | | betadisper | | |
|---|---|---|---|---|---|---|---|
| **comparison** | ***df*** | ***F*** | ***p**** | **$R^2$** | ***df*** | ***F*** | ***p*** |
| shrub – trees | 1 | 5.2303 | 0.001 | 0.1921 | 1 | 3.8377 | 0.06291 |
| conifer trees – broadleaf trees | 1 | 2.5358 | 0.048 | 0.1298 | 1 | 0.1969 | 0.6628 |

*df*, degrees of freedom; *F,* F-statistic; *p*, p-value; * Bonferroni adjusted p-value.

**Table S10** Standardized effect size (SES) of mean pairwise distance (MPD) calculated for all communities (samples), without taking into account species abundance, and using shuffled tip labels of a pruned tree as a null model (1000 iterations on 999 runs). The negative *z*-scores (mpd.obs.z) indicate that most *Methylobacterium* communities were composed of phylogenetically closely related species.

| sample | ntaxa | mpd.obs | mpd.rand.mean | mpd.rand.sd | mpd.obs.rank | mpd.obs.z | mpd.obs.p |
|---|---|---|---|---|---|---|---|
| ABBA_M11 | 3 | 0.1684 | 0.3603 | 0.0842 | 20 | -2.2786 | 0.02 |
| ABBA_M12 | 9 | 0.2525 | 0.3587 | 0.034 | 15 | -3.12 | 0.015 |
| ABBA_M13 | 5 | 0.2552 | 0.3618 | 0.0552 | 53 | -1.9322 | 0.053 |
| ABBA_M14 | 9 | 0.2454 | 0.3603 | 0.0339 | 7 | -3.3915 | 0.007 |
| ABBA_M15 | 6 | 0.1791 | 0.3608 | 0.0483 | 6 | -3.7614 | 0.006 |
| ACSA_M21 | 3 | 0.3208 | 0.3595 | 0.0886 | 285 | -0.4361 | 0.285 |
| ACSA_M22 | 2 | 0.3532 | 0.356 | 0.1374 | 405 | -0.0206 | 0.405 |
| ACSA_M24 | 5 | 0.2534 | 0.3589 | 0.0554 | 55 | -1.9042 | 0.055 |
| ACSA_M25 | 10 | 0.2159 | 0.36 | 0.0303 | 1 | -4.7547 | 0.001 |
| COCO_M01 | 14 | 0.308 | 0.3591 | 0.0206 | 19 | -2.4838 | 0.019 |
| COCO_M02 | 14 | 0.3179 | 0.3592 | 0.0205 | 32 | -2.0208 | 0.032 |
| COCO_M03 | 14 | 0.2415 | 0.3599 | 0.0213 | 1 | -5.5689 | 0.001 |
| COCO_M04 | 7 | 0.229 | 0.3607 | 0.0424 | 15 | -3.1112 | 0.015 |
| COCO_M05 | 19 | 0.3679 | 0.358 | 0.012 | 778 | 0.8299 | 0.778 |
| FAGR_M16 | 11 | 0.2857 | 0.3598 | 0.0275 | 17 | -2.6902 | 0.017 |
| FAGR_M17 | 9 | 0.2465 | 0.3605 | 0.0342 | 9 | -3.3303 | 0.009 |
| FAGR_M18 | 1 | *NA* | *NA* | *NA* | *NA* | *NA* | *NA* |
| FAGR_M19 | 6 | 0.2335 | 0.3592 | 0.0465 | 17 | -2.7054 | 0.017 |
| FAGR_M20 | 5 | 0.2552 | 0.3618 | 0.0552 | 53 | -1.9322 | 0.053 |
| THOC_M06 | 6 | 0.2446 | 0.3616 | 0.0463 | 32 | -2.5267 | 0.032 |
| THOC_M07 | 8 | 0.2591 | 0.3603 | 0.0382 | 22 | -2.6465 | 0.022 |
| THOC_M08 | 10 | 0.2467 | 0.3599 | 0.031 | 7 | -3.6573 | 0.007 |
| THOC_M09 | 12 | 0.2831 | 0.3586 | 0.026 | 10 | -2.9034 | 0.01 |
| THOC_M10 | 10 | 0.243 | 0.36 | 0.031 | 6 | -3.7734 | 0.006 |

ntaxa, number of *Methylobacterium* species in sample; mpd.obs, observed mean pairwise distance (mpd) between species within a community; mpd.rand.mean, mean mpd in null communities; mpd.rand.sd, standard deviation of mpd in null communities; mpd.obs.rank, rank of observed mpd compared to null communities; mpd.obs.z, standardized effect size of mpd compared to null communities: negative z-scores indicate phylogenetic clustering, while positive z-scores indicate larger distance than expected by chance; mpd.obs.p, *p*-value of observed mpd compared to null communities.

**Table S11** Comparison of gene clusters composition of *Methylobacterium* communities between *C. cornuta* and all four tree host species, and between conifer and broadleaf trees, using PERMANOVAs and multivariate homogeneity of variances tests (betadisper).

| | PERMANOVA | | | | betadisper | | |
|---|---|---|---|---|---|---|---|
| **comparison** | ***df*** | ***F*** | ***p****** | **$R^2$** | ***df*** | ***F*** | ***p*** |
| shrub – trees | 1 | 3.0322 | 0.004 | 0.12617 | 1 | 11.562 | 0.0027 |
| conifer trees – broadleaf trees | 1 | 2.0041 | 0.006 | 0.11131 | 1 | 0.7255 | 0.4069 |

*df*, degrees of freedom; *F,* F-statistic; *p*, *p*-value; * Bonferroni adjusted *p*-value.

### 3 – SUPPLEMENTARY FIGURES

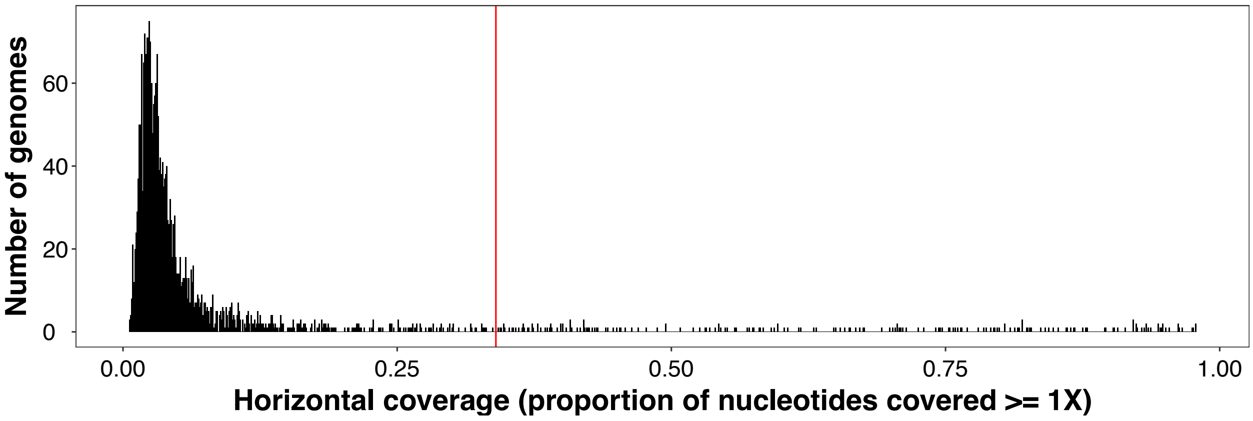


(A)

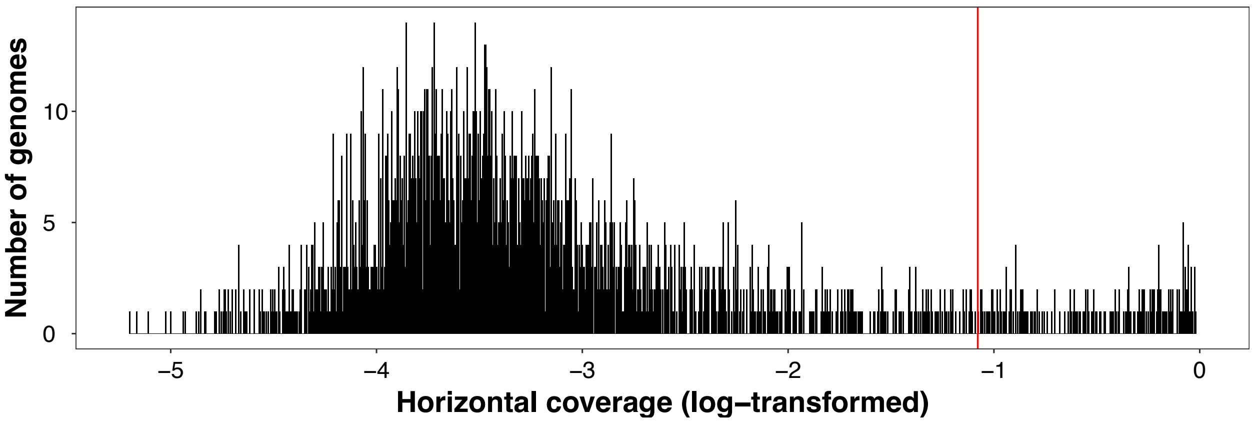


(B)

**Figure S1** Distribution histograms of horizontal coverage values for all 104 genomes in all 25 samples ($n$ = 2600). Log-normal distribution (A) and log-transformed normal distribution (B). Red lines represent values with a $z$-score of 1.96 calculated on the log-transformed normal distribution.

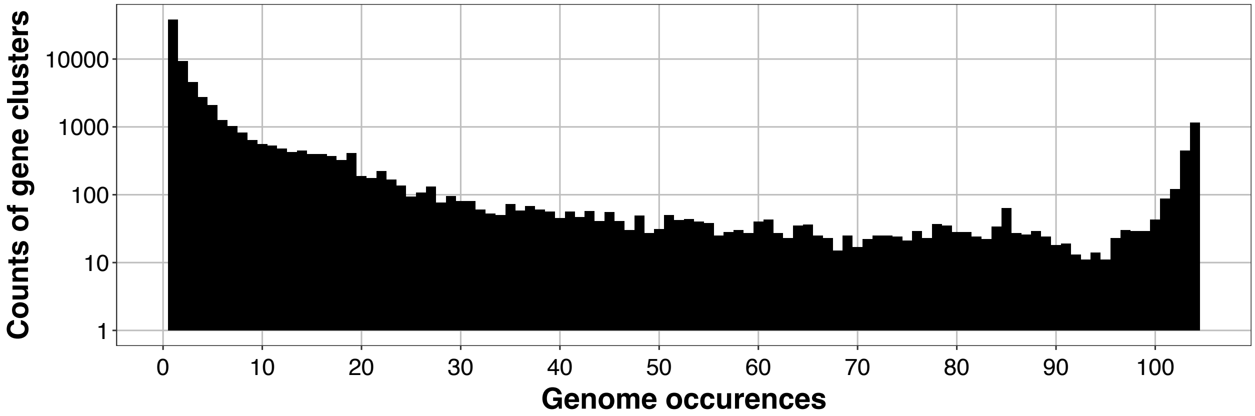


**Figure S2** Distribution of gene cluster counts by genome occurrences within the pangenome of *Methylobacterium*. This figure illustrates how many different gene clusters are present in a given number of genomes. For example, around 1000 gene clusters are found in all 104 genomes composing the pangenome.

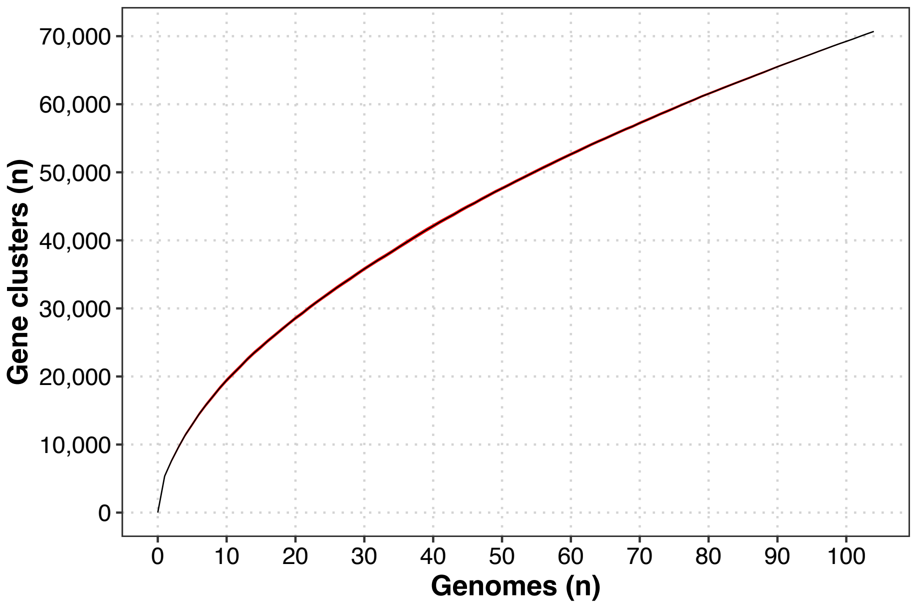


**Figure S3** *Methylobacterium* had an open pangenome pattern: the number of new gene clusters increased continuously with increasing number of genomes considered in the pangenome, without reaching a plateau. The curve shows a power regression following Heaps law (Heap's law test, $\alpha = 0.376$). We used the R package *micropan* [3] to analyze the *Methylobacterium* pangenome openness. We used the function 'rarefaction' (100 permutations) to generate the rarefaction curve illustrating the cumulative number of distinct gene clusters in the pangenome with increasing genome number. Black line represents the mean values of permutations, and the red area, the 95% confidence interval. We used the function 'heaps' (100 permutations) to test if the pangenome was open ($\alpha < 1$) or closed ($\alpha > 1$), according to Tettelin et al. [4] who suggested that an open pangenome follows a Heaps' law [5]. Heaps test's result was $\alpha = 0.376$.

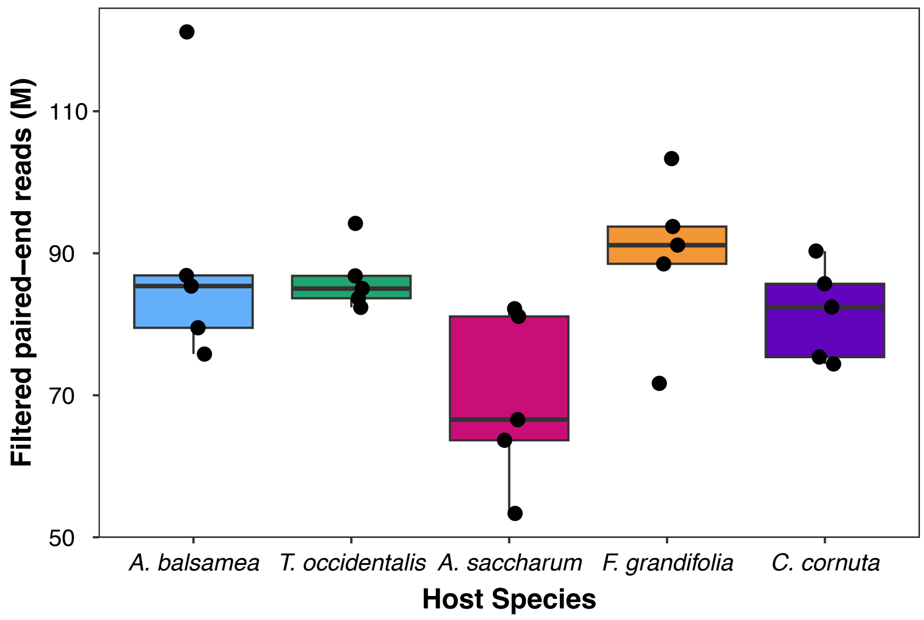


**Figure S4** The number of filtered paired-end reads (in millions) obtained from metagenomic shotgun sequencing did not significantly differ among the five host species (ANOVA, $p = 0.063$, adj.$R^2 = 0.217$), although there was a tendency for *A. saccharum* samples to yield fewer reads.

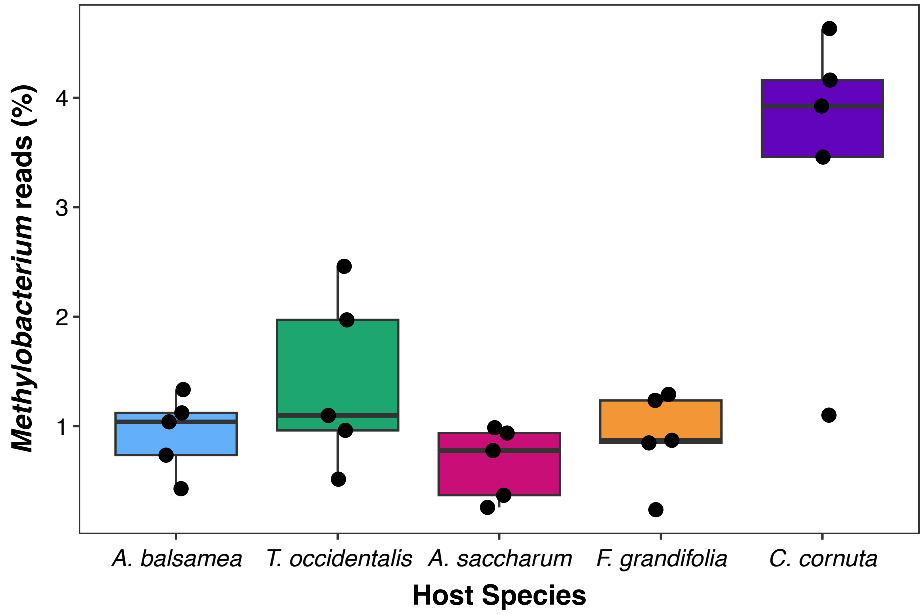


**Figure S5** The metagenomes of *C. cornuta* were generally composed of a larger proportion of *Methylobacterium* sequences than the metagenomes of other host species (Kruskal-Wallis, $p = 0.021$, $\eta^2 = 0.377$). The boxplot illustrates the percentage of metagenomic single reads that mapped to *Methylobacterium*'s pangenome in samples of the five host species.

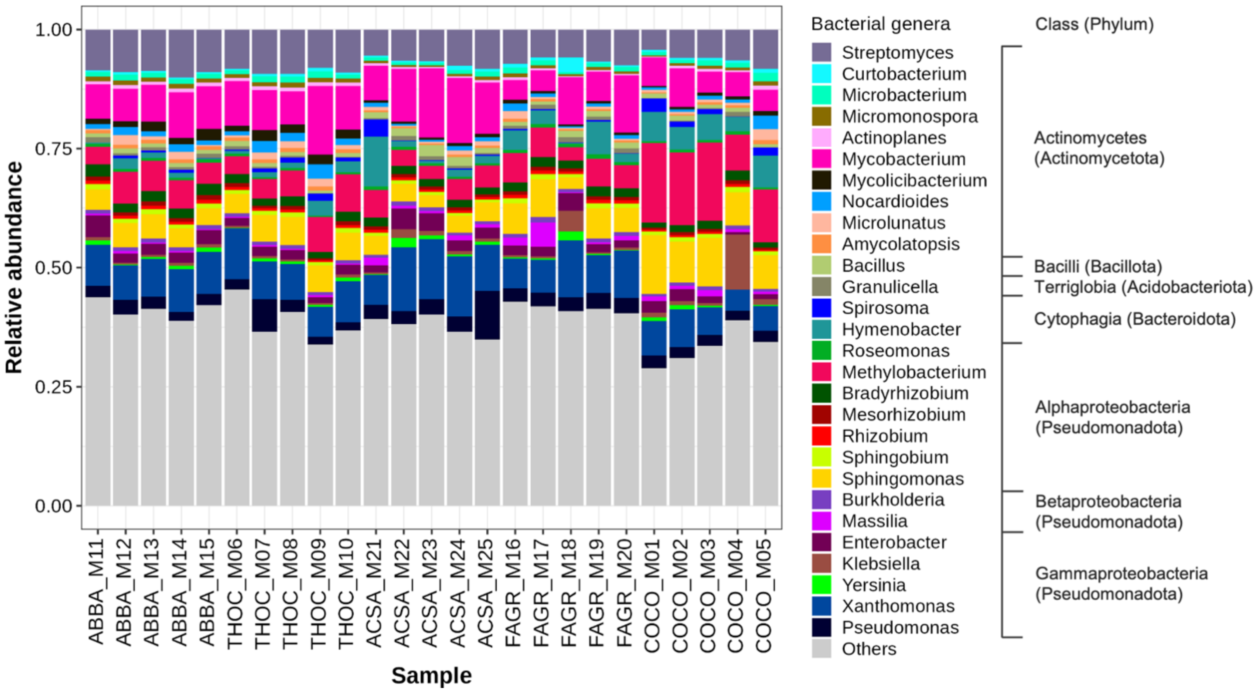


**Figure S6** The proportion of bacterial genera composing the phyllosphere communities varied among the 25 samples and among the five host species. The barplot illustrates the 28 most abundant genera, ordered by class and phylum. “Others” category includes all genera whose reads did not account for at least 0.5% of the total bacterial reads across samples. The most prevalent bacterial genera across all samples were *Mycobacterium* (8.42 ± 3.09%), *Xanthomonas* (8.41 ± 2.37%), *Streptomyces* (7.49 ± 1.48%), and *Methylobacterium* (6.77 ± 3.98%).

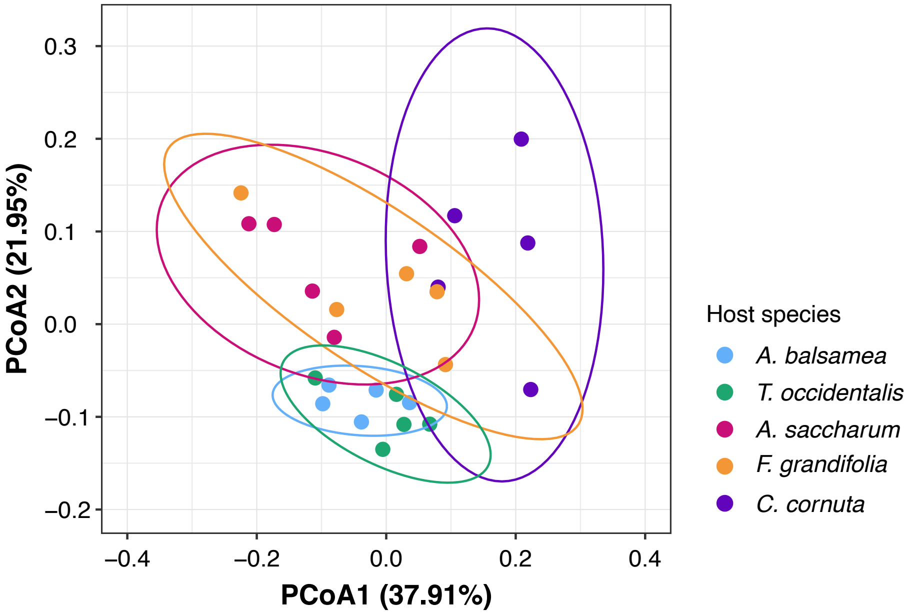


**Figure S7** Genus-level community composition differed between *C. cornuta* and tree species, as well as between conifer and broadleaf tree species (PERMANOVAs, $p$ = 0.001) (Table S7). The first axis of the PCoA highlights the distinction between C. cornuta and other host species communities. The second axis highlights the distinction in composition between conifers and broadleaf host species. The relatively smaller distance among conifer communities compared to the broadleaf communities reflect their higher homogeneity in genus composition (Table S7). Bray-Curtis dissimilarity coefficient was used to calculate distances between pairs of samples. Ellipses indicate 95% confidence intervals.

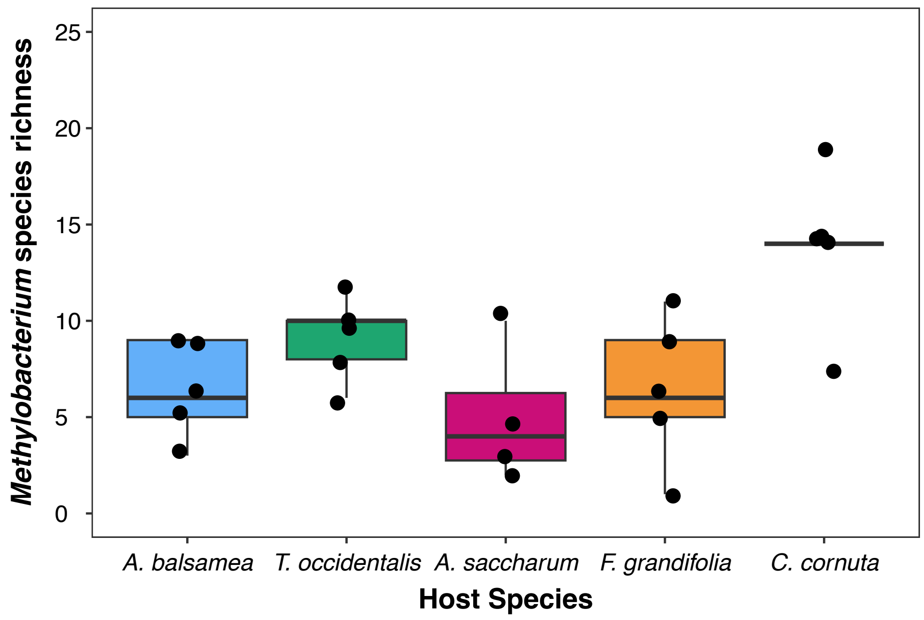


**Figure S8** *Methylobacterium*'s community species richness was higher in *C. cornuta* samples (Table S8, $p$ = 0.040). The shrub species harboured in average over 13 *Methylobacterium* species, while tree species generally harboured around 7 species. However, the Shannon index calculated for Methylobacterium communities did not vary with host species (KW, $p$ = 0.336, $\eta^2$ = 0.029).

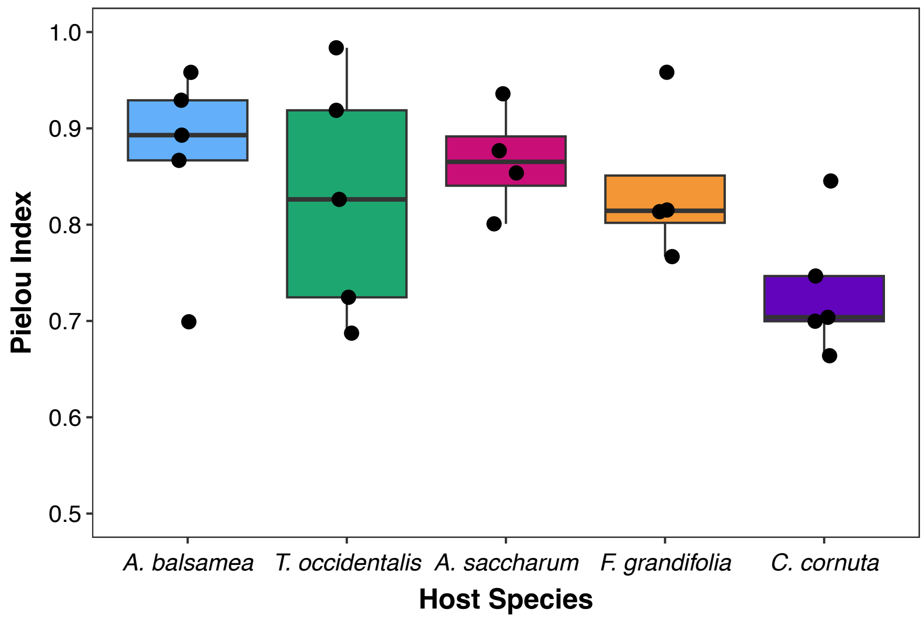


**Figure S9** *Methylobacterium*'s community evenness, as measured by the Pielou index, did not significantly differ among the five host species (ANOVA, $p$ = 0.176. adj.$R^2$ = 0.125), although *C. cornuta* communities, which are richer in species, tended to be slightly less even.

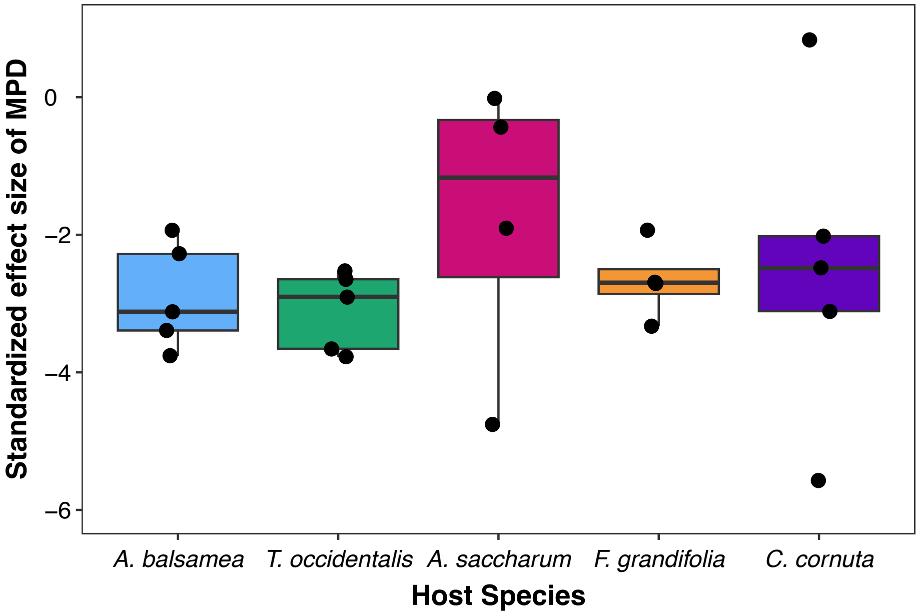


**Figure S10** The standardized effect size of mean pairwise distance ($SES_{MPD}$) – a measure of community phylogenetic dispersion expressed as *z*-scores – of *Methylobacterium* species within communities did not vary among the five host species (ANOVA, $p = 0.729$).
